# ProSolo: Accurate Variant Calling from Single Cell DNA Sequencing Data

**DOI:** 10.1101/2020.04.27.064071

**Authors:** David Lähnemann, Johannes Köster, Ute Fischer, Arndt Borkhardt, Alice C. McHardy, Alexander Schönhuth

## Abstract

Obtaining accurate mutational profiles from single cell DNA is essential for the analysis of genomic cell-to-cell heterogeneity at the finest level of resolution. However, sequencing libraries suitable for genotyping require whole genome amplification, which introduces allelic bias and copy errors. As a result, single cell DNA sequencing data violates the assumptions of variant callers developed for bulk sequencing, which when applied to single cells generate significant numbers of false positives and false negatives. Only dedicated models accounting for amplification bias and errors will be able to provide more accurate calls.

We present ProSolo, a probabilistic model for calling single nucleotide variants from multiple displacement amplified single cell DNA sequencing data. It introduces a mechanistically motivated empirical model of amplification bias that improves the quantification of genotyping uncertainty. To account for amplification errors, it jointly models the single cell sample with a bulk sequencing sample from the same cell population—also enabling a biologically relevant imputation of missing genotypes for the single cell. Through these innovations, ProSolo achieves substantially higher performance in calling and genotyping single nucleotide variants in single cells in comparison to all state-of-the-art tools. Moreover, ProSolo implements the first approach to control the false discovery rate reliably and flexibly; not only for single nucleotide variant calls, but also for artefacts of single cell methodology that one may wish to identify, such as allele dropout.

ProSolo’s model is implemented into a flexible framework, encouraging extensions. The source code and usage instructions are available at: https://github.com/prosolo/prosolo

## 1 Introduction

Originally, genome sequences have been queried for genetic germline variation or for highly abundant somatic variation, for example in cancer. The advent of high-throughput single cell sequencing has recently turned the spotlight on a type of so far understudied variation: the often less abundant somatic or post-zygotic variation that constantly accumulates with every mitotic cell division throughout the lifetime of an organism, turning every individual into a complicated genomic mosaic^1^. Estimates for somatic single nucleotide variants (SNVs) range from around 0.6 · 10^−9^ up to 60 · 10^−9^ mutations per genome position per cell division^2–5^, with a recent estimate^6^ based on single-cell sequencing at 2.66 · 10^−9^. With the size of the human (reference) genome at approximately 3.2 · 10^9^ base pairs, these numbers indicate that even during healthy development, most cells harbour cell-specific point mutations. This enables retrospective monitoring of lineages involved in normal organism development, merely by sampling some cells^7^, without having to interfere with its general development, or kill the individual. In other words, this establishes a universally applicable methodology for *in vivo* lineage tracing.

In cancer development, this variation can be used to trace the cellular ancestry of tumour subclones and metastases^8–11^, and to characterise the evolutionary dynamics of cancer progression^12, 13^. In the long run, methods that account for the dynamics of mutational signatures in cellular evolution will improve diagnosis, treatment and prognosis of diseases for which somatic alterations are a key factor. To this end, obtaining accurate profiles of the genetic variation affecting single cells is essential.

In sequencing libraries prepared directly from single cells, only a small fraction of the genome is sampled. To obtain coverage levels that allow for the consistent identification of SNVs across larger parts of the genome, in *vitro* whole genome amplification is crucial. Among whole genome amplification methods, multiple displacement amplification (MDA14) has proven the least error-prone and is therefore considered the state-of-the-art in single cell SNV profiling^15–18^. But, although the type of polymerase used in MDA (Φ29) has the highest fidelity currently attainable (due to its proof-reading functionality), amplification errors still occur at a rate of 1.24 · 10^−6^ to 9.5 · 10^−6^ per copied base^15, 19–22^—three orders of magnitude higher than the estimates for the somatic mutation rate. Further, the efficiency and fidelity of Φ29 polymerase depends on the template sequence context^23^, implying that the amplification error rate systematically varies around this average. Moreover, the degree of amplification depends on the quality of the template DNA extracted from the single cell^24^ and how accessible each stretch of DNA is to amplification initiation via priming^25^. As a result, sequencing coverage after amplification differs both between sites along the genome and between the two alleles at a particular site, up to the entire dropout of alleles^26^. Because standard variant callers assume that alleles are uniformly covered, they do not perform well on the resulting data and are substantially outperformed by single-cell variant callers^27, 28^. Clearly, when calling SNVs for single cells, the statistical uncertainties introduced by the amplification need to be dealt with at the largest possible accuracy.

Thus, variant callers for whole-genome-amplified single-cell data need to account for both amplification errors and allelic bias, in addition to accounting for the site-specific variation. However, state of the art single cell SNV callers routinely assume fixed global rates when modeling uneven allelic coverage (up to dropout) and amplification errors. For example, to reflect the amplification error rate, both MonoVar^27^ and SCcaller^28^ work with global false positive error rates for calling the presence of an alternative allele at a particular site. This assumes that Φ29 polymerase is agnostic to local template sequence context, although it is not^23^. Similarly, for modelling allele dropout, MonoVar^27^ and SCIPhI^29^ assume that one rate applies globally (and across all cells). This neglects that allele dropout, as the extreme case of uneven allele coverage, varies greatly along the genome and in particular also between cells, because their DNA is amplified separately. Interestingly, SCIPhI^29^ additionally models allelic amplification bias to be governed by one global beta-binomial distribution (to apply for all cells in a dataset), thereby accounting for allelic dropout a second time (as the extreme values at 0 and 1 of that distribution). Improving on that point, tools exist that account for local variation in allelic amplification bias. SCcaller^28^ and SCAN-SNV^30^ both estimate the minor allele frequency from nearby germline heterozygous sites. However, SCcaller employs a fixed global false positive error rate for the calling of alternative alleles^28^, and SCAN-SNV makes use of heuristics for filtering candidate variants^30^. Thus, to the best of our knowledge, there is no statistical model that allows for local variation of bias and errors due to amplification, and for statistically sound false discovery control when calling and genotyping SNVs in single cells.

We describe ProSolo, a variant caller using a unifying statistical framework that takes into account all relevant MDA related biases and errors, allowing for them to vary locally. Importantly, our model further enables a computationally efficient implementation, which is challenging even in bulk variant calling when considering local effects due to statistical uncertainties affecting the data^31^. ProSolo’s statistical rigor allows for an accurate control of the false discovery rate when calling alternative alleles or identifying other relevant effects, such as allele dropout.

## 2 Results

### 2.1 Single Cell Sequencing Model

We describe a novel probabilistic model that addresses the genotyping of diploid single cells whose DNA has been subject to whole genome multiple displacement amplification (MDA)^14^. In the following, we introduce the central innovations of our model and demonstrate its advantages in comparison to existing approaches. More details and a detailed derivation of all model elements can be found in the Supplement, including a summary of the core model.

Our model addresses the two major issues of MDA: (i) the differential amplification of the two alleles present in a diploid cell (”amplification bias” in the following); (ii) MDA induced errors (”amplification errors” in the following) which are copy errors introduced by the Φ29 polymerase used in MDA. To address amplification bias, we leverage a mechanistically motivated, empirically derived model of differential amplification of alleles. To assess amplification errors, we evaluate single cell samples together with a bulk sample from which the single cell is supposed to stem. Regarding the latter, we argue that a bulk sample should be added to single cell sequencing experiments wherever possible: it samples from the same cell population without requiring amplification, and is therefore unaffected by amplification bias and errors and thus makes a particularly useful background sample to address the statistical uncertainties and biases induced by MDA. At the same time, one of the major features of the core model and its implementation is that it can easily be adapted to flexibly deal with other sampling setups, so it could be extended to further scenarios. For related work on flexible bulk sequencing sample composition, see Köster et al.^31^.

#### A mechanistically motivated model of amplification bias, trained on data, gives realistic coverage-specific single-cell (genotype) likelihoods

To account for MDA amplification bias up to the complete dropout of individual alleles, we distinguish between two alternative allele frequencies: (i) The theoretical underlying allele frequency at a site in a single cell: *θ_s_*. This can be assigned one of three possible values, namely *θ_s_* ∈ {0,0.5,1}, where 0 and 1 represent the homozygous reference and alternative genotype and 0.5 a heterozygous genotype. However, the ratio of reads harboring the different alleles from a single cell sequencing experiment does not reflect the true allele frequency, because of the biases induced by amplification. Instead, the ratio of reads reflects (ii) the allele frequency after its distortion through amplification bias. For a site with total coverage of *l* reads, of which *k* reads bear the alternative allele, the formal definition of this measurable frequency is 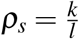. To accurately quantify the uncertainty introduced by amplification bias, we consider

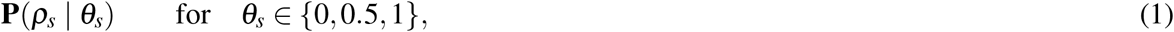

the probability distributions that reflect the shift from the true allele frequency *θ_s_* to the distorted allele frequency *ρ_s_*, as induced by MDA (Figure 1C). We thus formally describe the statistics of read counts skewed by MDA at all sites, encompassing sites that are homozygous for the reference allele, heterozygous, or homozygous for the alternative allele.

**Figure 1.**
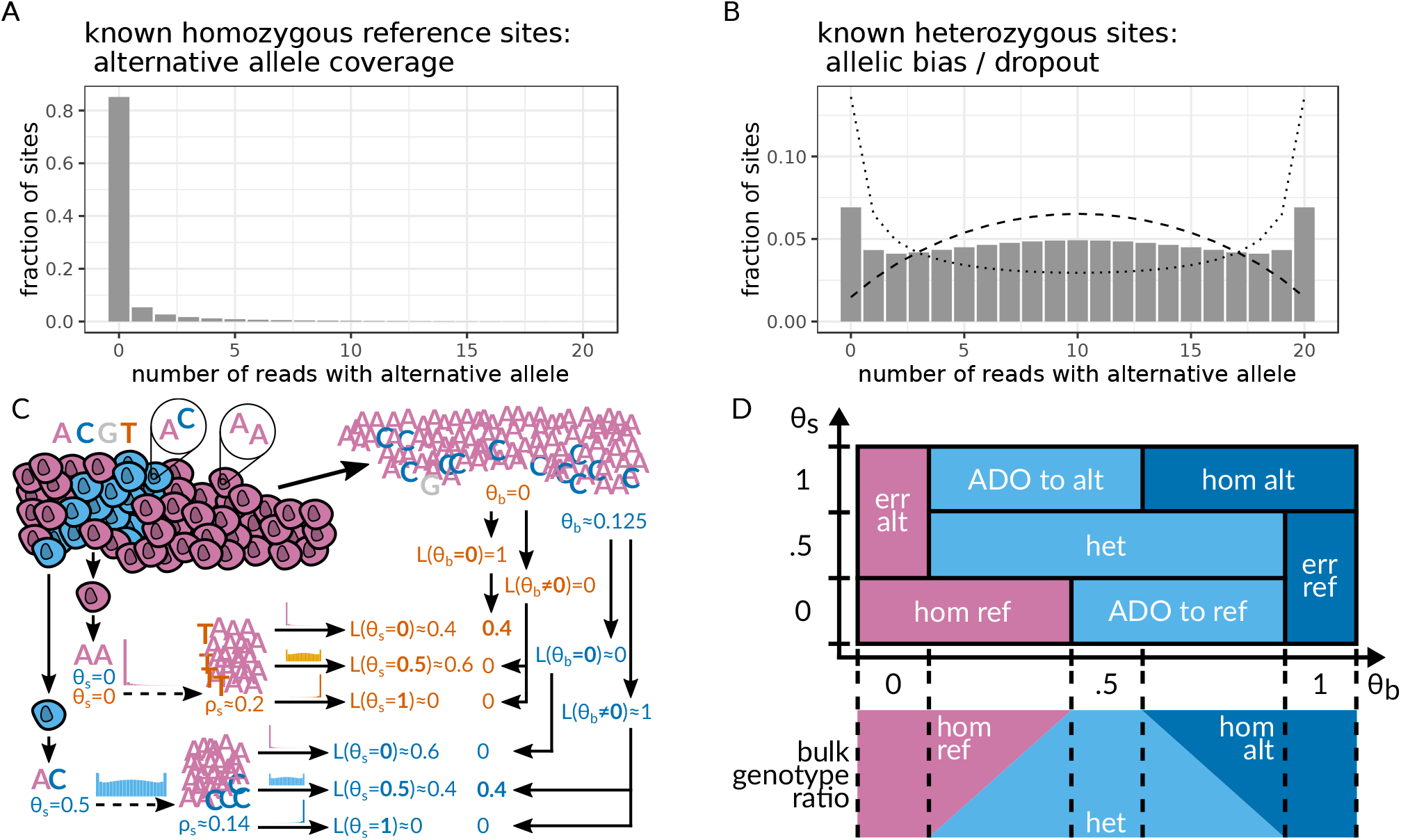
**A,B** Exemplary alternative allele read count distributions for sites covered by 20 reads, as derived by Lodato et al.22 Homozygous reference sites in **(A)** are assumed to follow a beta-binomial distribution; sites heterozygous for the alternative allele in **(B)** are assumed to follow the linear combination of two symmetrical beta-binomial distributions (dotted and dashed lines). **(C)** Toy example of calling the same genomic site in two single cells from the same population that differ in their true underlying allele frequencies for alternative allele C (blue, *θ_s_* = 0 vs. *θ_s_* = 0.5). Alternative nucleotide T (orange) is an amplification error. Empirical distributions in A and B account for the amplification bias, and likelihoods for the alternative allele candidates from the bulk reduce the likelihoods of amplification errors, thereby correctly identifying both the error and the original true mutation. This is formalised with the model in D. **(D)** Definition of single cell events based on true underlying alternative allele frequencies in the single cell (*θ_s_*) and the bulk (*θ_b_*) (assuming that the bulk sample has a deep coverage that captures somatic variants). The bulk is always assumed to be a combination of a maximum of two genotypes at a particular site, generating all possible *θ_b_* (bottom panel). ADO – allele dropout, alt – alternative, err – error, het – heterozygous, hom – homozygous, ref – reference.

To do this, we follow the considerations of Lodato et al.^22^ (see Figure S5 and the section “Modeling MDA-derived alternative read counts” of the respective supplement for the details), who fitted well-studied probability distributions to empirical distributions they obtained for **P**(*ρ_s_* | *θ_s_*) of *θ_s_* ∈ {0,0.5}. For sites that are homozygous for the reference allele, amplification bias cannot initially happen. However, once an amplification error creates an alternative allele, this can be amplified to large frequencies due to amplification bias. Lodato et al.22 thus consider the empirical distribution **P**(*ρ_s_* | *θ_s_* = 0) to follow a betabinomial. Effectively, this means that the probability of a non-zero alternative read count (*ρ_s_* > 0, which is any alternative read count above 0 in Figure 1A) will be non-zero (**P** (*ρ_s_* | *θ_s_* = 0) > 0), merely because of sequencing and amplification errors. In contrast, at heterozygous sites (Figure 1B), the distribution is dominated by amplification bias. Thus, for **P**(*ρ_s_* | *θ_s_* = 0.5) they found a mixture of two beta-binomial distributions to appropriately fit the empirically observed distributions (Figure 1B). We further motivate the choice of the beta-binomial distribution mechanistically by an analogy to its generative urn model named after Pólya^32^: Take an urn with two white and two black balls, where each ball represents one strand of each (double-stranded) allele at a heterozygous site. In the Pólya urn model, drawing a ball leads to replacement of that ball with two balls of the same color, analogous to a strand copy by the Φ29 polymerase (additional discussion in the Supplement, Section S 1.2.1). Finally, the necessity for a mixture of beta-binomials (Figure 1B) becomes evident when contrasting it with the binomial distribution observed at heterozygous sites in bulk experiments. Namely, the bulk distribution for reads supporting the alternative allele would narrowly peak at a count of 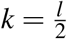, which is not the case for the corresponding *k* = 10 in Figure 1B. Instead, the mixture of beta-binomials peaks at the extreme read counts of *k* = 0 and *k* = *l*, highlighting that the dropout of the alternative or reference allele is quite likely to occur. Most likely, this mixture arises from differences in the accessibility of different pieces of the genomic DNA for amplification^25^ (Supplement, Section S 1.2.1). Finally, for homozygous alternative sites (i.e. **P**(*ρ_s_* | *θ_s_* = 1)), we rely on the symmetry of the cases *θ_s_* = 0 and *θ_s_* = 1. In summary, we obtain the following equations for **P**(*ρ_s_* | *θ_s_*):

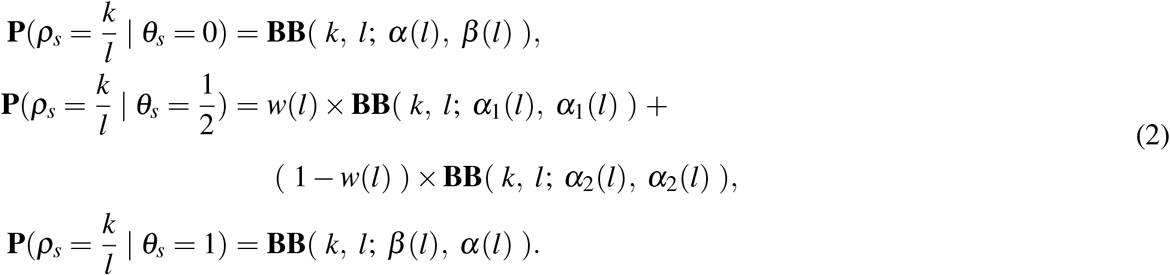

Here, **BB** represents the beta-binomial probability mass function where *k* ∈ {0, …, *l*} with *l* being the total read coverage of the site that is considered. All of parameters *α, α*_1_, *α*_2_, *β*, and *w* scale linearly in *l*. Allowing to vary distributions through these parameters allows amplification bias to depend on the total coverage of a site, and thereby to vary locally. Symmetry of distributions for the homozygous sites (*θ_s_* = 0 in Figure 1A, and *θ_s_* = 1) is established by swapping the shape parameters *α* (*l*) and *β* (*l*). The distribution for heterozygous sites (Figure 1B) corresponds to a mixture of two beta-binomials with shape parameters *α*_1_(*l*) = *β*_2_(*l*) and *α*_2_(*l*) = *β*_2_(*l*), where equality of *α*(*l*) and *β*(*l*) yields symmetry of the distributions relative to 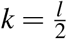 (the expected value for alternative read counts at heterozygous sites). Slopes and intersects for the scaling of all parameter values across different choices of *l* are given in Table S 1.

#### Using bulk evidence of alternative alleles to increase the accuracy of variant calls

A bulk sequencing sample of the same cell type from the same organism is a much larger sampling of that cell population than the sequencing of dozens of single cells. Unless a single cell is the only one to harbour a particular mutation, a deep enough sequencing of the bulk sample from which the single cell was drawn should contain reads from cells that share the particular mutation with the single cell. Thus, considering bulk samples corresponds to drawing unbiased samples of the entire population (of genome copies), in contrast to single cells that correspond to heavily distorted measurements on pairs of copies. This establishes a formal, statistical argument to why one should consider bulk experiments in single cell sequencing whenever possible and why the identification of a mutation in an accompanying bulk sample lends further credibility to the mutation in the single cell in a data-driven way, without assuming any fixed error rate (Figure 1C). As a consequence, bulk samples can be employed to improve both the sensitivity and specificity of variant calls in the single cell, where increasing depth of coverage of the bulk sample increases the accuracy of the calls.

For our model, we derive likelihoods for all possible alternative allele frequencies in the background bulk sample. Given a set of *n* reads from the bulk (*b*) read data 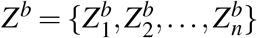, and discrete possible allele frequencies 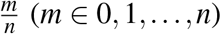, we compute the probability of the data given a particular allele frequency as the product of the probabilities of all the reads:

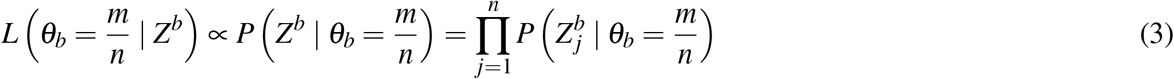

Here, the probability of an individual read, given a particular allele frequency, 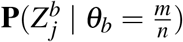, is defined as in Equation S 21, based on the model described in Köster et al.^31^.

#### Calculating posterior probabilities for events at single cell sites, including a bulk background sample

With single cell genotype likelihoods adjusted by our empirical amplification bias model (Equation 2) and auxiliary evidence on alternative alleles from a bulk sample (Equation 3), we define mutually exclusive single cell events. Figure 1C gives a simplified illustration of how the combination of likelihoods works for calling genotypes in the single cell. However, our model fully defines the two-dimensional space of possible underlying alternative allele frequencies in the two samples as:

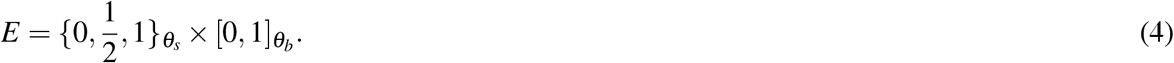

Within this space, we define single cell events as mutually exclusive subspaces, for example an error-free homozygous alternative site is defined by allele frequencies 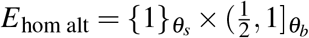 or the dropout of an alternative allele across allele frequencies 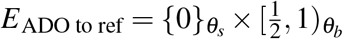 (Figure 1D, Table S 2). We thus obtain a set of mutually exclusive single cell events (Figure 1D):

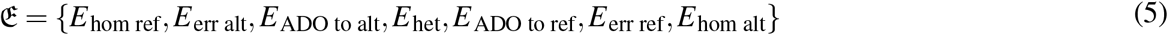

Assuming a flat prior across the possible underlying allele frequencies for both the bulk and the single cell, we can compute likelihoods for all those single cell events (e.g. Equation S 28). The sum of the likelihoods of all these (mutually exclusive) events yields the marginal probability (Equation S 26). Using the marginal probability, we can calculate the posterior probability for any of these events. For example, the posterior probability of event *E*_hom alt_ (Figure 1D) can be calculated with:

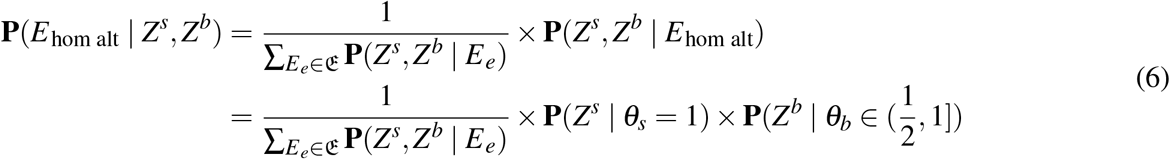

Accounting for the sample likelihoods based on Equation S 23 (assuming *ρ_b_ = θ_b_* for the bulk that has no amplification step, Equation S 3), and evaluating only point estimates of these likelihoods at possible alternative allele frequencies, this gives:

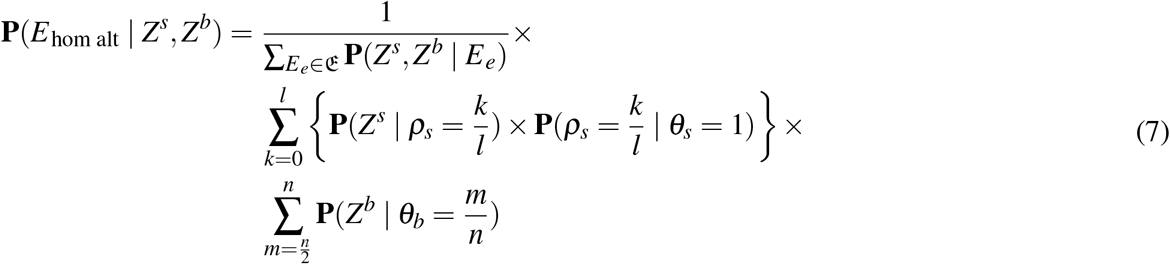

Accounting for amplification bias with Equation 2 and computing the likelihood of the sample-specific allele frequency ranges with Equations S 22 and 3, this becomes:

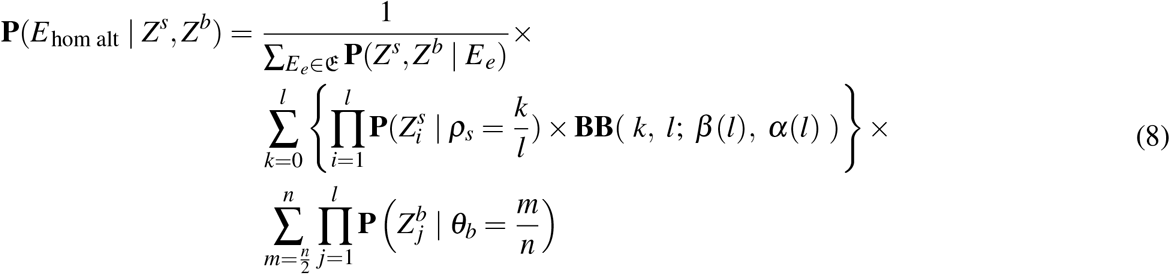

With read probabilities calculated with Equation S 21, we can calculate the posterior probability of the *E*_hom alt_ event (for an analogous and more detailed derivation for *E*_ADO to ref_, see Supplement, Section S 1.4). To obtain posterior probabilities for compound events, we sum up the posterior probabilities of the events it comprises, effectively joining up their respective allele frequency range combinations into compound range combinations. For the site-specific single cell probability for the presence of an alternative allele in the single cell, we thus get the compound event (blue events in Figure 1D; Table S 2):

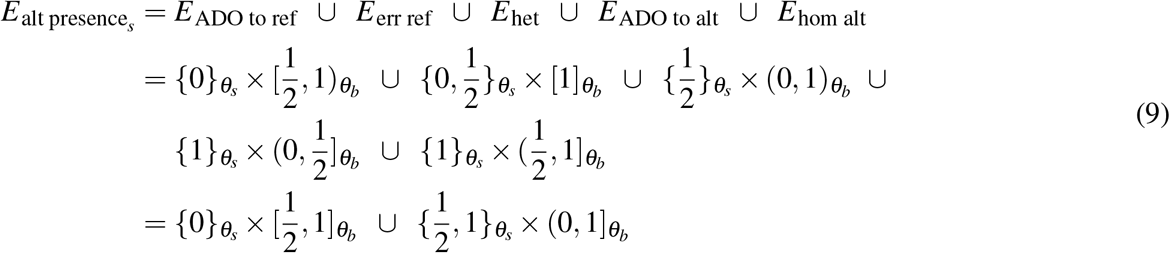

whose posterior probability we can obtain from this sum:

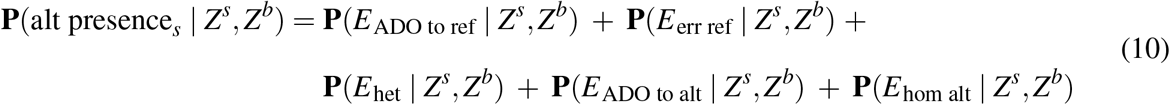

To genotype, we calculate the posterior probability of all three possible single-cell genotypes and choose the genotype with the maximum posterior probability. In Figure 1D, events are colored purple when implying the homozygous reference genotype, light blue when implying the heterozygous genotype and dark blue when implying the homozygous alternative genotype (Table S 2). Similarly, to calculate the posterior probability of an allele dropout at a particular site, we sum up the posterior probabilities of the two `ADO` events.

For any such compound event, we can estimate a threshold on the posterior probabilities that controls for a specified false discovery rate. This is based on the approach described by Müller et al.^33, 34^—for further details see the Supplement (Section S 1.6) and Köster et al^31^.

#### Biologically relevant imputation based on the bulk sample

As argued above, a large enough sampling of the bulk cell population that the single cell comes from should contain the single cell’s genotype at a particular site, unless this cell is genuinely the first cell to harbour a mutation at that site. This bulk background sample can thereby render credibility to single cell variants with low coverage, while at the same time eliminating amplification errors in the single cell sample, as these will not exist in the bulk sample. Interestingly, the bulk sample also provides a mechanism of biologically meaningful imputation at sites that have no read coverage in the single cell. If imputation is desired for sites with no read coverage in the single cell, we set 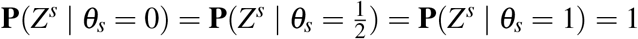, rendering all (unknown underlying) single cell genotypes equally likely. Thus, the posterior probabilities of events at sites with no read coverage become solely dependent on the read data from the bulk sample, providing the most common genotype in the bulk. However, while this is a biologically meaningful way of imputation at the vast majority of genomic sites, it should be noted that this imputation will usually favor germline genotypes over any existing (lower frequency) somatic genotypes at a site.

#### ProSolo is an easy-to-use command-line tool, based on a modular framework

ProSolo is an easy-to-use command-line tool—following usability standards^35^—and its source code is available at https://github.com/prosolo/prosolo, including instructions for an easy installation via Bioconda^36^. Its main contribution in terms of software is the implementation of its comprehensive statistical model into the Rust variant calling library of Varlociraptor^31^. See the Supplement (Section S 1.6) for further implementation details.

### 2.2 Benchmarking

We compare ProSolo to the state-of-the-art for SNV calling from single cell sequencing data of multiple displacement amplified (MDA) DNA: MonoVar^27^, SCAN-SNV^30^, SCcaller^28^ and SCIPhI^29^. We used Snakemake^37^ (version `5.4.0`) to implement the benchmarking workflows. For detailed information on benchmarking setup and results, see Supplementary Section S 2. All code used for benchmarking is available at: https://github.com/prosolo/benchmarking_prosolo (or as preserved by Zenodo at: https://doi.org/10.5281/zenodo.3769116).

#### 2.2.1 Datasets and Ground Truths

We benchmarked ProSolo on two experimental datasets (Figure 2), each with a different kind of ground truth:

**Figure 2.**
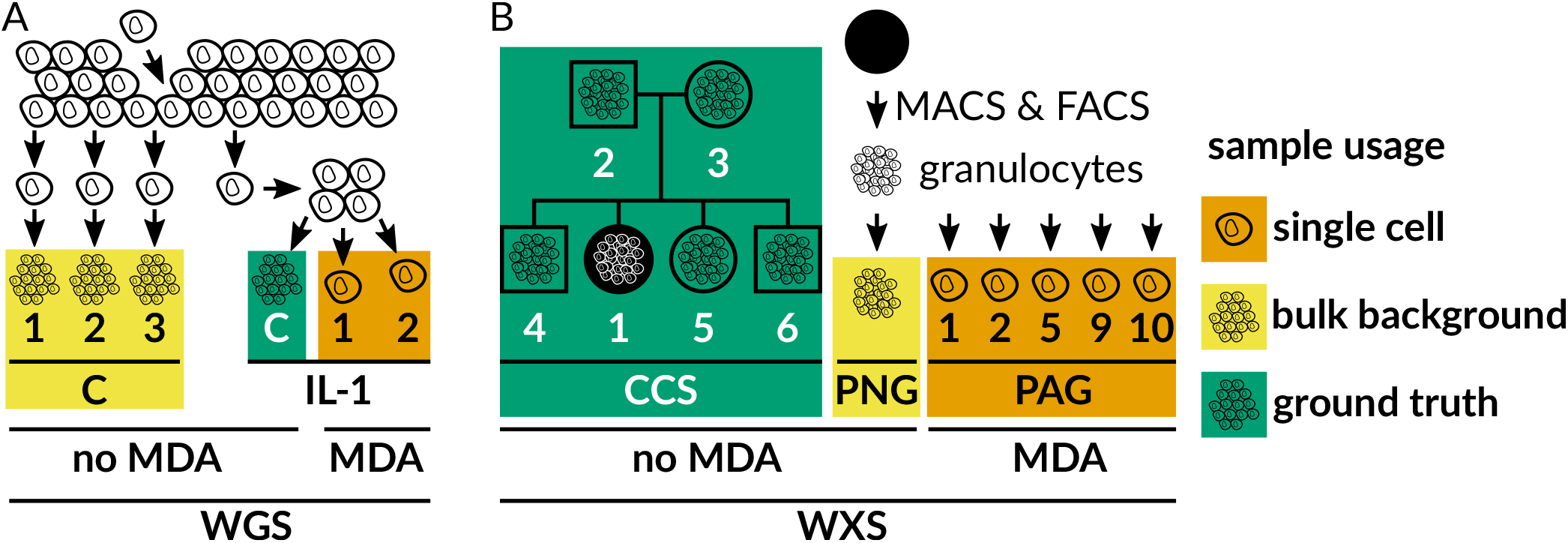
Benchmark Datasets. For single cells, DNA was multiple displacement amplified (MDA). **A** Whole genome sequencing (WGS) dataset, generated from a clonal population started with an individual cell^28^, and expanded further downstream to generate different bulk samples (C1, C2, C3, IL-1C). **B** Newly generated whole exome sequencing (WXS) dataset of blood cells. Ground truth genotypes of patient CCS1 were determined from sequencing data of family members (boxes male; circles female). Granulocytes were isolated from blood using magnetic and fluorescence activated cell sorting (MACS and FACS).

##### Whole genome sequencing of almost identical kin-cells from a cell line^28^

The first dataset comes from the publication of the SCcaller software^28^ (Figure 2A, dataset available from project PRJNA305211, accessions SRR2976561 to SRR2976566). A single starting cell was grown in two steps (Figure 1 of the original paper28): After an initial mini-expansion, a single cell was selected as the founder for the secondary IL expansion into 20-30 cells. From this, two cells were extracted (IL-11 and IL-12) and sequenced following MDA. The remaining kindred cells from that clone were used as a bulk sequencing sample without amplification (IL-1C). IL-1C serves as the ground truth, as these cells are only very few cell divisions away from IL-11 and IL-12, and thus have almost no difference in the somatic mutations acquired. The ground truth genotype was generated using GATK HaplotypeCaller to call variant sites and bcftools mpileup to identify homozygous reference sites (with read coverage above 25 but no alternative allele present). IL-1C was only used as a ground truth and not provided as input to any of the software compared here. Three more distant clones (C1, C2, C3), generated from cells after the first mini-expansion, were merged into a further bulk sample for SCcaller and ProSolo (see Software and Parameters below). Unlike other callers (all of which finished in less than 5 days), SCIPhI took five weeks to finish on this dataset in sensitive mode and 7.5 weeks in default mode.

##### Whole exome sequencing of five human granulocytes with a pedigree ground truth

For the second benchmarking dataset, blood was taken from a patient with a constitutional mismatch repair-deficiency_38_ after informed consent. Granulocytes were selected via Magnetic-Activated Cell Sorting (MACS) and Fluorescence Activated Cell Sorting (FACS, Figure 2B, dataset available as EGAD00001005929 in EGA study EGAS00001004123). Individual cells were isolated using a microfluidics device and subjected to multiple displacement amplification (MDA). Using a panel of 16 primer pairs covering different genes across chromosomes for quantitative real-time PCR, we selected granulocytes where at least 15 of these loci were properly amplified. For those cells, we performed whole exome capture, sequencing library preparation and paired-end Illumina sequencing. From the remaining sorted cell population, we also extracted bulk DNA and submitted it to whole exome capture and paired-end Illumina sequencing without MDA, to generate a bulk background sample for ProSolo and SCcaller.

To generate the ground truth of this dataset, we could leverage previously sequenced bulk whole exome data from the same person, their parents and three siblings^38^ (Figure 2B). To create ground truth germline alternative allele calls, we ran three pedigree-aware variant callers (BEAGLE4.0^39^, polymutt^40^ and FamSeq^41, 42^) and created a consensus by including only calls where all callers agree at a site and where a maximum of one caller not calling the site was allowed (Figure S 4).

#### 2.2.2 ProSolo achieves highest alternative allele calling accuracy

The most precise single cell variant callers to date, SCcaller and SCIPhI, only call the presence vs. the absence of an alternative allele (i.e. the heterozygous and the homozygous alternative genotypes called jointly, the joint probability of the blue event areas in Figure 1D). We thus focused on this for the main benchmarking.

The whole genome cell line dataset (Figure 2A) seems much less challenging than the other dataset, as all methods achieved very high precision in alternative allele calling (Figures 3A and S 5A), at recall rates of 45% and higher. In comparison to all other tools, ProSolo shows striking increases in recall. For example, an increase of nearly 10% for a precision above 99%, where its maximum recall is 76.6%, compared to 70.5% for MonoVar, 68.7% for SCIPhI and 61.0% for SCcaller. The only exception with a recall of 0.01% is SCAN-SNV (at a precision of 99.2%). This can be explained by it aiming at somatic mutations, while the vast majority of SNVs in a genome will be germline variants.

**Figure 3.**
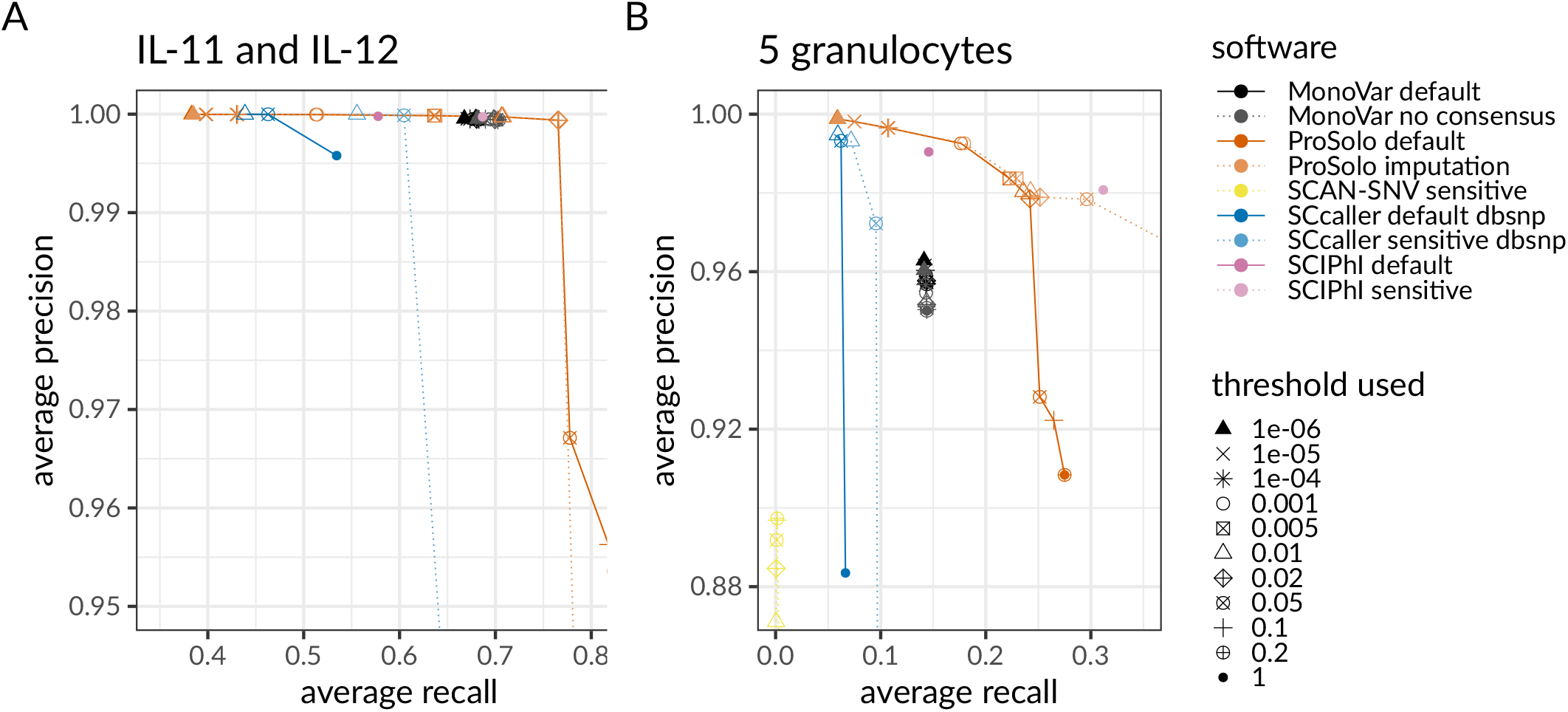
Precision-recall plots for alternative allele calls of ProSolo, MonoVar, SCAN-SNV, SCcaller and SCIPhI. Note that both panels are strong zoom-ins, focusing on (different) areas of interest. Global views of these panels are provided in Figure S 5. **A** Precision and recall average of two whole genome sequenced single cells IL-11 and IL-12 against their kindred clone IL-1C as ground truth genotypes. **B** Precision and recall average of the five whole exome sequenced single granulocytes against their pedigree-based germline genotype ground truth. Threshold parameters: MonoVar `--t`; ProSolo `--fdr`; SCAN-SNV `--fdr`; SCcaller `-a cutoff`; SCIPhI `none available`. Software modes: MonoVar with consensus filtering (*default*) or without (*no consensus*); ProSolo with minimum coverage 1 in single cell (*default)*, or imputing zero coverage sites based on bulk sample (*imputation*); SCcaller with recommended settings (*default*) or with a more *sensitive* calling; SCIPhI with default parameters (*default*) or all heuristics off (sensitive).

Although a relative increase in recall of about 10% at utmost precision is certainly remarkable, ProSolo demonstrates its power on the second (whole exome) dataset (Figure 2B). For this dataset, only SCcaller, SCIPhI and ProSolo achieved a precision above 99%, with ProSolo reaching a 20% increase of recall to 17.6%, compared to SCIPhI’s 14.6%, and SCcaller with 7.2% (Figures 3B and S 5B). In comparison, MonoVar achieved a maximum precision of only 96.29%. However, this was at a much higher recall (14.13%) than for example SCcaller (9.55% at a precision of 97.23%). SCcaller’s decreased recall on this dataset might be due to its estimation of local allelic bias by also taking biases at neighboring sites into account—in whole exome data the number of neighboring sites available for this estimation will be limited and might lead to less reliable estimates.

On this dataset, SCAN-SNV’s recall increased to 0.16% at a decreased maximum precision of 89.7%. Most likely, this decreased precision is an artefact of using the germline genotype as ground truth. At the sites with somatic mutations in single cells, which SCAN-SNV focuses on, this ground truth will instead contain the homozygous reference germline genotype and will incorrectly classify (existing) alternative alleles as false positives. Due to this effect, we also expect the calculated precision of all the other tools to be an underestimate. However, as the other tools also provide alternative allele calls for all sites where the single cells retained this germline genotype, the relative effect on their precision will be smaller.

At the same time, this germline ground truth caveat also indicates that the recall of SCIPhI’s sensitive mode and ProSolo’s imputation mode will be an overestimate. Whenever coverage of a site is missing in a single cell, SCIPhI may impute the genotype with the last common ancestor genotype of the most closely related cells, while ProSolo will impute to the majority genotype in the bulk sample. Both strategies provide a biologically meaningful imputation that will be more useful than post-hoc modes of imputation. However, at single cell sites where a somatic mutation has created a true alternative allele, but no coverage is provided, we expect that both methods are most likely to call the homozygous reference germline genotype. In then comparing this to the germline genotype as the ground truth, these calls will be classified as true negatives even though they really constitute false negatives, thus artificially increasing recall. While the underestimation of precision equally affects all tools and generally means that benchmarking results are more conservative than with a more accurate (somatic) ground truth, this overestimation of recall in only these modes of two tools does not allow for a fair comparison. We have thus excluded both SCIPhI’s sensitive mode and ProSolo’s imputation mode from the discussion of the whole exome dataset with its germline ground truth (but their results are nevertheless displayed in Figures 3B, S 5B, S 6B and S 7B for reference).

Finally, a feature where ProSolo clearly stands out is the control over the false discovery rate. As can be seen in Figure 3 (and Figure S 5), ProSolo provides flexible control over precision vs. recall via specifying a false discovery rate of interest. While no other tool provides a formal control over the easily interpretable false discovery rate, several of the tools provide other types of thresholds that we varied in attempts to achieve higher precision or recall. However, none of them provide control over similar ranges of precision and recall. The only limit to that range with ProSolo’s current model, is that it becomes less accurate when controlling for very small false discovery rates (below 0.01% for alternative allele calling in the whole genome dataset, see Supplement Section S 2.4). But this still leaves ProSolo as the only tool that provides the user with the choice of either aiming for more discoveries at the cost of a higher rate of false discoveries, or at aiming for a more limited number of discoveries with higher confidence in each of them.

#### 2.2.3 Estimates of allele dropout rate validate ProSolo’s model

Leveraging our ground truths and using three different ways to calculate the allele dropout rate, we can confirm the general validity of our single-cell event definitions and also explore limitations of the current model. For the allele dropout rate, we will focus on the set of sites where the respective ground truth call is heterozygous, as these are the sites where the dropout of one of the alleles can be identified in a non-ambiguous manner.

More details for the three ways in which we calculate allele dropout rates are given in the Supplement (Section S 2.5). Here, we give a short explanation:

1. At each ground truth heterozygous site, we sum the posterior probabilities of the two allele dropout events defined in ProSolo (”ADO to alt” and “ADO to ref” in Figure 1D) to obtain a total allele dropout probability (Equation S 31) and use these to compute the expected value of allele dropout across all sites. This gives us an expected allele dropout rate (”expected value” in Figure 4, Equation S 34). 2. We genotype all the ground truth heterozygous sites with ProSolo, take the most likely genotype and then compare against the ground truth: heterozygous sites that ProSolo calls as homozygous are counted as dropout sites. The number of such sites is divided by the total number of ground truth heterozygous sites where the respective single cell had coverage (”hom at ground truth het” in Figure 4, Equation S 35). 3. We identify all heterozygous ground truth sites with a coverage of at least 7, where without amplification bias we could be reasonably sure to sample both alleles (for the reasoning see Supplement, Section S 2.5). We then count a site as an allele dropout if there is one allele (reference or alternative) with no read coverage at all, and again divide by the total number of ground truth heterozygous sites where the respective single cell had coverage (”no alt/ref read at ground truth het” in Figure 4, Equation S 36).

**Figure 4.**
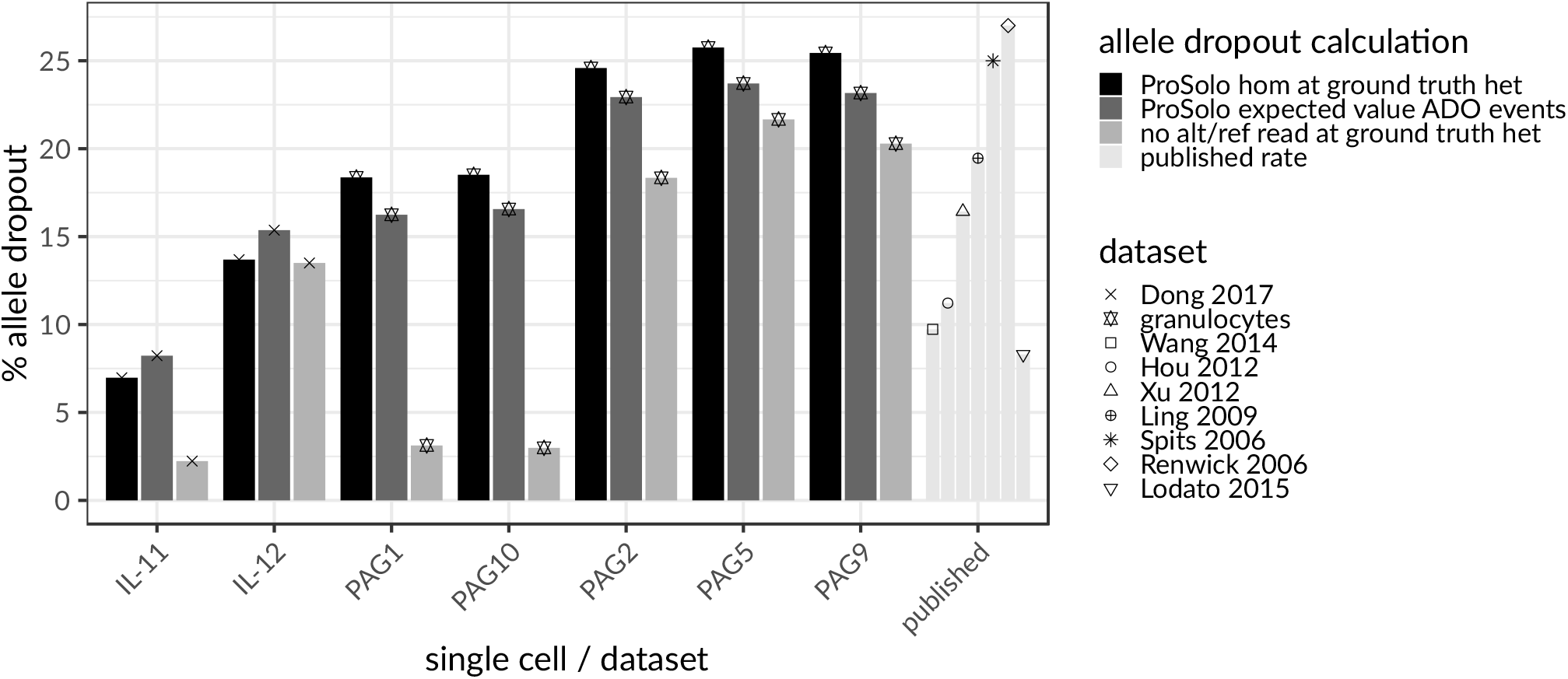
Concordance of three differently calculated per-cell (IL-1 and PAG cells) allele dropout rates across ground truth heterozygous sites, in the context of allele dropout rates from the literature^21, 22, 26, 43–47^. The expected value of the allele dropout events in ProSolo (dark gray) is concordant with the number of false homozygous genotype calls made by ProSolo on those sites (black) and both values are well within the range of published allele dropout rates for single cell MDA sequencing data (”published”, to the right, very light gray). The naive allele dropout rate (light gray)—calculated as ground truth heterozygous sites with a minimum coverage of seven and either no read with the reference allele or no read with the alternative allele—shows discrepancies with ProSolo’s estimates of allele dropout for samples with a more uniform coverage (IL-11, PAG1, PAG10, Figure S 3).

The expected allele dropout rates based on the ProSolo probabilities for allele dropout clearly fall into the range of previously published allele dropout rates^21, 22, 26, 43–47^ (”published” in Figure 4). This analysis also clearly shows that the ProSolo expected allele dropout rates, based on the model’s probabilities, correspond to those determined by comparing ProSolo genotypes with the ground truth (Figure 4). This demonstrates that the explicit modelling of allele dropout events works and is useful for genotyping. However, in that comparison, the expected allele dropout rate was consistently underestimated on our own whole exome data (”granulocytes”, Figure 4), and slightly overestimated for the data from Dong et al.^28^ (”Dong 2017”, Figure 4). The comparison to a naively calculated allele dropout rate consistently shows an overestimation of the allele dropout probability, which is strongest for the samples with a higher overall coverage (IL-11, PAG1, PAG10, Figures 4 and S 3). However, this overestimation of the allele dropout probability does not seem to impact the genotyping resolution (Figures S 8 and S 9).

## 3 Discussion

ProSolo is the first method for SNV calling from MDA single cell sequencing data to comprehensively model both amplification bias and amplification errors in a way that allows for site-specific variation (Figure 1). We achieve this by combining a data-driven model of amplification errors that incorporates a bulk background sample alongside the single cell sample with an empirical model of amplification bias, based on a statistical understanding of the MDA process that is mechanistically motivated and dependent upon a site’s coverage. The underlying model calculates posterior probabilities for fine-grained single cell event definitions, whose false discovery rate can be controlled for and that can pass more information about data uncertainties to probabilistic models in downstream analyses. Using a whole genome and a whole exome dataset—each with a different type of ground truth (Figure 2)—we demonstrate that these two innovations of ProSolo combined to achieve better accuracy (Figure 3). Moreover, the model allows to accurately control the false discovery rate of variant calls, another novelty of great practical value, and both the model and its modular implementation are easy to adapt and extend.

Especially the joint modeling of a single cell and a bulk sample is favorable from a statistical point of view: in terms of MDA induced bias, the bulk acts as an unbiased sample of the population from which the single cell was drawn. This provides a drastically more sensitive approach compared to, for example, the consensus rule of calling alternative alleles only when there is evidence in at least two (or even three) single cells (as implemented in MonoVar^27^). A more systematic and biologically relevant model of sharing information is implemented in SCIPhI, where the phylogenetic relationship of cells is part of the estimated parameters. Intuitively, if two cells are closely related, they have a higher likelihood of sharing a genotype at a particular site. However, both the consensus rule and the sharing of information via an inferred phylogeny requires that more than one single cell exhibit sufficient coverage of an alternative allele to call it reliably. This will rarely be achieved in single cell MDA experiments, which are often limited to a few dozen cells. In contrast, bulk sequencing can easily be scaled to a much larger sampling of a cell population—representing hundreds or thousands of cells of a cell population—simply by adding a single higher coverage sample. Our benchmarking demonstrated that ProSolo achieves a substantially higher recall than any other tool for a precision above 99% in identifying the presence of an alternative allele. To summarize our benchmarking of alternative allele calling: SCcaller seems optimized for precision, MonoVar achieved higher recall than SCcaller, and SCAN-SNV does not seem suitable for general variant calling on MDA single cell sequencing data, but is only applicable when restricting interest to somatic variants. ProSolo clearly achieves the best accuracy, with particularly striking increases in recall. The only other tool that shows consistent performance across both datasets, but with a recall 10 to 20% below ProSolo, is SCIPhI. However, SCIPhI takes several weeks on a single core without any possibility of parallelization, compared to a runtime of only a few days on a cluster for all the other tools.

Jointly calling single cell variants with a bulk background sample also enables a biologically relevant imputation of genotypes at sites where single cells lack coverage. If genotype profiles without missing values are for example required by downstream tools, ProSolo can provide a cell-population-specific imputation instead of resorting to a majority assignment (limited by the number of single cells sequenced) or even imputation based on external databases (e.g. dbsnp). As our results further demonstrate, ProSolo’s empirical model of amplification bias is robust also for whole exome sequencing data. In contrast, SCcaller—that estimates local amplification bias based on minor allele frequencies at known germline genotypes in the vicinity of an examined site—suffers from a scarcity of neighbouring sites in whole exome data.

Finally, ProSolo is the only method that provides a clearly interpretable false discovery rate parameter that can actually control the trade-off between precision and recall. This can be used on alternative allele calls (Figure 3, Supplement Section S 2.4), the main focus of our benchmarking, but also on any of the events pertinent to single cell analysis (such as allele dropout or amplification errors) or combinations thereof (see Figure 1D for all events). Thereby, beyond just calling alternative allele presence in a statistically reliable way, ProSolo can, for example, also compute the expected allele dropout rate across the entire genome of a particular cell in a robust manner (Equation S 31, Figure 4).

In addition to such global rates, ProSolo determines reliable site-specific posterior probabilities for all single cell events that might be of interest, including allele-resolved genotypes, allele dropout or amplification errors. We anticipate that such fine-grained probabilities—and the uncertainties they capture—will be informative for improving probabilistic modelling in downstream analyses. They could e.g. prove useful in models for phylogenetic reconstruction of the lineage relationship of sequenced single cells^29, 48, 49^, while keeping them computationally tractable^13^.

An in-depth look at one of the above-mentioned fine-grained events, allele dropout, showed that ProSolo’s allele dropout rate estimates were within the range of published estimates. This confirms that our modeling of allele dropout events is realistic and can be useful in alternative allele calling and genotyping. However, the allele dropout rate estimations were slightly off in different directions for the different benchmarking datasets, suggesting that the empirical distributions we currently make use of (based on Lodato et al.22) may not suit all datasets. For example, whole genome amplification may introduce further variability so far not captured by our model. This conclusion is further supported by the naively calculated allele dropout rate, which is much lower in the high coverage cells of the two datasets (IL-11 for the Dong et al. data^28^, PAG1 and PAG10 in our granulocytes; see Figure S 3). Interestingly, the sample IL-11 from Dong et al.28 has been suggested to be a doublet^30^, which might explain the higher overall coverage and points to a possible source for the discrepancy between the naively calculated and the estimated allele dropout rates. If samples PAG1 and PAG10 were doublets as well, this would indicate that our use of empirical distributions in ProSolo provides for more robust event probabilities in the presence of doublets, while heuristic thresholding (as in our naive allele dropout estimate) is very sensitive to such perturbations. In general, the use of a fixed empirical model for MDA allelic bias does not seem to impede ProSolo’s performance in alternative allele calling compared to the other tools, but has a noticeable effect on genotyping (Supplement, Section S 2.6) and might be responsible for slight imprecisions when controlling for very small false discovery rates (Supplement Section S 2.4). When future datasets are generated based on improved MDA protocols^13^, these effects might be exacerbated. We therefore consider it important future work to improve on the fitting of the empirical read count distributions observed when applying MDA to single cells (Supplement, Section S 1.2.2). For example, the parameters of the mechanistically motivated combination of beta-binomial distributions for modelling heterozygous genotypes could be learned per single cell sample at germline heterozygous sites—similar to the approaches of SCcaller and SCAN-SNV, but globally per cell with their local variation modelled by their dependence on a site’s coverage. Further room for improvement also remains for the modeling of the homozygous genotypes: while the current distributions account for both amplification and sequencing errors, sequencing errors are already safely accounted for elsewhere in our latent variable model. An amplification error profile that is not compounded with sequencing errors, would need to be based on the **Φ**29 polymerase error rate and a better understanding of the statistical distribution that these errors generate (Supplement, Section S 1.2.2). Studying and implementing the corresponding changes in the future has the potential to further improve the accurate site-specific event probabilities that ProSolo already provides through the joint modelling with a bulk background sample.

Finally, the modular implementation of ProSolo within the context of the Varlociraptor library^31^ facilitates the implementation of further features, such as the ones mentioned above, read-backed phasing^50, 51^ or even more variable models of amplification bias^30^ that could be integrated to exhaustively leverage MDA data information content. Since the Varlociraptor library also provides advanced functionality for the calling of insertions, deletions and multiple nucleotide variants (MNVs), one of the immediately following next steps will be to adapt ProSolo to calling those in single cells.

Overall, ProSolo provides an accurate and easy-to-use variant caller for single cell MDA sequencing data, which will empower more research using single cell sequencing data.

## Supporting information

Supplementary Methods and Results

## 4 Online Methods

See Supplement PDF.

