## Supplementary Methods and Results for "ProSolo: Accurate Variant Calling from Single Cell DNA Sequencing Data"

David Lähnemann, Johannes Köster, Ute Fischer,

Arndt Borkhardt, Alice McHardy, Alexander Schönhuth

April 27, 2020

### S 1 Probabilistic model of ProSolo

#### S 1.1 Underlying True Alternative Nucleotide Frequencies: $\theta$

The eventual goal in variant calling is to infer the true frequency of a possible alternative, and thus variant, nucleotide in the sample you sequenced. For such an alternative nucleotide $x \in \{A, C, G, T\}$ at a particular site in a genome, we therefor define[1]:

$$\theta_b(x), \theta_s(x) \tag{S 1}$$

as the (true, but usually unknown) frequency of alternative nucleotide $x$ in the bulk and single cell sample, respectively. These frequencies translate into probabilities of sampling a DNA fragment containing the respective nucleotide at a site: either directly, as with the sequencing of DNA from a bulk of cells $b$; or indirectly, as with the sequencing of whole genome amplified DNA of a single cell $s$. It is our eventual goal to work back from bulk and single cell data to derive likelihoods for possible values of $\theta_b(x)$ and $\theta_s(x)$, respectively, allowing us to place likelihoods on possible underlying genotypes and events like allele dropouts in single cells.

[1] We omit an explicit notation for a site, but every $\theta$ is always specific to one genome position.

For a single diploid cell $s$, the possible alternative nucleotide frequencies are

$$\theta_s(x) = \begin{cases} 0 \\ \frac{1}{2} \\ 1 \end{cases} \qquad \text{(S 2)}$$

reflecting the cases that: (i) $x$ is not present at the position—if no other allele is present, this constitutes a homozygous reference site; (ii) $x$ reflects an alternative nucleotide that is present in only one of the two genome copies of the single cell—this constitutes a heterozgyous site; (iii) $x$ is an alternative nucleotide present in both genome copies of the single cell—this constitutes a homozygous alternative site.

Bulk experiments contain subpopulations that can differ in their genotype regarding a particular site, through a somatic mutation that one of the subpopulations has acquired in one allele. They can thus exhibit any alternative nucleotide frequency ($\theta_b(x) \in [0, 1]$), depending on the genotypes present and their relative abundance (Figure S 2).

#### S 1.2 Non-reference Nucleotide Frequencies After Biased Whole Genome Amplification: $\rho$

Let further $\rho_s(x), \rho_b(x)$ be the abundance of genome copies showing alternative nucleotide $x$ at a particular site in the single or bulk DNA sample, respectively, right before the sequencing—for the single cell, this means after the whole genome amplification that is necessary to obtain enough DNA for sequencing with technologies like Illumina. These abundances translate into probabilities to draw a read containing $x$ from the respective sample (in the sequencing experiment), given that such a read covers the genome position.

Under the assumption of uniform abundance of all parts of the genome and uniform abundance of each haplotype, which is reasonable in bulk sequencing (i.e. without amplification), it holds that $\rho = \theta$, so we henceforth assume that

$$\rho_b(x) = \theta_b(x). \tag{S 3}$$

In contrast, in single cell sequencing—due to the necessary whole genome amplification process—the assumption of uniform abundance of all parts of the genome and of each haplotype does not hold. Instead, each allele at a particular site can be amplified to a different degree, ranging from no amplification at all (i.e. a complete allele dropout) to several hundred or thousand copies. Thus

$$\rho_s(x) \neq \theta_s(x) \quad \text{but instead} \quad \rho_s(x) \sim P(\rho_s \mid \theta_s(x)) \tag{S 4}$$

that is, $\rho_s$ is a random variable governed by a probability distribution $P(\rho_s \mid \theta_s(x))$ that depends on the original alternative nucleotide frequency $\theta_s$. $P(\rho_s \mid \theta_s(x))$ reflects the differential amplification of the two alleles, which is only detectable for heterozygous sites ($\theta_s = \frac{1}{2}$), thus

$$P(\rho_s \mid \theta_s(x)) = \begin{cases} P(\rho_s \mid \theta_s(x) = 0) \\ P(\rho_s \mid \theta_s(x) = \frac{1}{2}) \\ P(\rho_s \mid \theta_s(x) = 1) \end{cases} \tag{S 5}$$

Understanding these probability densities is key to understanding the statistical issues single cell whole genome amplification methods come along with[2]. From the Supplementary Material of Lodato et al. [2015] (see Figure S5 and the section "Modeling MDA-derived alternative read counts" of the respective supplement for the details), we get an empirical model that characterizes them for the whole genome amplification method

[2] Note, in addition to the whole genome amplification bias, these distributions can implicitly also model the amplification errors that the polymerases used for whole genome amplification introduce in the copy process.

multiple displacement amplification (MDA)[3]:

$$
\begin{aligned}
P(\rho_s \mid \theta_s = 0) &= \mathfrak{BB}_{l,\alpha,\beta}(\rho_s) \\
P(\rho_s \mid \theta_s = \frac{1}{2}) &= w(l) \times \mathfrak{BB}_{l,\alpha_1,\alpha_1}(\rho_s) + (1 - w(l)) \times \mathfrak{BB}_{l,\alpha_2,\alpha_2}(\rho_s) \\
P(\rho_s \mid \theta_s = 1) &= \mathfrak{BB}_{l,\beta,\alpha}(\rho_s)
\end{aligned}
\tag{S 6}
$$

where $\mathfrak{BB}$ represents beta-binomial functions with the parameters linearly scaled by $l$, the total observations at a site (i.e. the total read coverage). For homozygous reference sites ($\theta_s = 0$), the parameters of the beta-binomial function are $\alpha(l)$ and $\beta(l)$. For the heterozygous sites ($\theta_s = \frac{1}{2}$), both beta-binomial functions are symmetrical with $\alpha_1(l), \alpha_2(l)$ dependent on the site-specific coverage $l$ and combined by factor $w(l)$. $\rho_s \in \frac{k}{l}, k \in 0, 1, \ldots, l$ are the possible cases of alternative nucleotide frequencies. In their Supplementary Figure S5 in panels C and D, Lodato et al. [2015] give estimates of the parameters $\alpha(l), \beta(l), w(l), \alpha_1(l), \alpha_2(l)$ obtained by fitting to empirical data. We list the values we extracted in Table S 1. For the third case (homozygous alternative nucleotide, i.e. $P(\rho_s \mid \theta_s = 1)$) we mirror the homozygous reference function[4].

Using one of the possible expressions of the respective probabilities (Equation 6.28 in Section 6.2.4 of Johnson et al. [2005])[5], with regards to the beta function $\mathfrak{B}$, we can

[3] For other whole genome amplification methods with different amplification bias properties, we can simply implement a different instance of the general model for $P(\rho_s \mid \theta_s(x))$ accordingly. And for MDA, different empirical models learned from other or more published datasets could be implemented as well as strategies of inferring parameters from the very dataset analysed in a certain run of the software.

[4] Note that the amount of sites that are homozygous for the reference allele is much larger, which also means that statistics for $\theta_s = 1.0$ are more robust when being derived from the case $\theta_s = 0.0$.

[5] Here, the parameters $b, w, c$ from the formulation in Equation 6.28 in Section 6.2.4 of Johnson et al. [2005] are summarised into $\alpha_1, \alpha_2$ from Lodato et al. [2015] with:

$$
\begin{aligned}
\alpha_1 &= \frac{w_1}{c} = w_1 = b_1 \\
\alpha_2 &= \frac{w_2}{c} = w_2 = b_2
\end{aligned}
\tag{S 7}
$$

I.e., the number of starting reference and alternative alleles is identical within each beta-binomial, yielding symmetric distributions. In terms of the Pólya urn model used as motivation in Johnson et al. [2005], this equates to the number of $w$ white and $b$ black balls being identical. $c$, the number

Table S 1: Parameters of the multiple displacement amplification (MDA) amplification bias model as presented in Supplementary Figure S5, panels C and D of Lodato et al. [2015]. Each of these scales linearly up to a site coverage of 60, with the slopes and intercepts as given here.

| parameter | slope | intersect |
|---|---|---|
| $\alpha$ | -0.000027183 | 0.068567471 |
| $\beta$ | 0.007454388 | 2.367486659 |
| $w$ | 0.000548761 | 0.540396786 |
| $\alpha_1$ | 0.057378844 | 0.669733191 |
| $\alpha_2$ | 0.003233912 | 0.399261625 |

compute the probability of observing $k$ reads with the alternative nucleotide at a site of coverage $l$ for the heterozygous case as follows:

$$\begin{aligned} P(K = k \mid \theta_s = \frac{1}{2}) = w(l) \times \binom{l}{k} \frac{\mathfrak{B}_{k+\alpha_1, l-k+\alpha_1}}{\mathfrak{B}_{\alpha_1,\alpha_1}} + \\ (1 - w(l)) \times \binom{l}{k} \frac{\mathfrak{B}_{k+\alpha_2, l-k+\alpha_2}}{\mathfrak{B}_{\alpha_2,\alpha_2}} \end{aligned} \quad \text{(S 8)}$$

In the future, these distributions could be learned from the dataset at hand, whenever large enough sets of sites that have a high probability of being homozygous reference or heterozygous are available. In addition, the exact distribution used could be further refined with the below considerations regarding the suitability of beta-binomial distributions in this case.

of balls of identical color that are added whenever a ball of particular color is added, is set to 1. Further considerations for obtaining the values of $b, w$ are discussed below in relation to the Pólya urn model.

##### S 1.2.1 The Pólya urn model: motivating the use of beta-binomial distributions

For the heterozygous case, the applicability of the beta-binomial distribution [Irwin, 1954] should be immediately evident from the formulation of the Pólya urn model [Eggenberger and Pólya, 1923] that generates beta-binomial distributions [Irwin, 1954, Johnson et al., 2005]. It works directly analogous to the amplification process[6]:

At the start, the urn contains $w$ *w*hite and $b$ *b*lack balls, analogous to the two alleles present at a heterozygous site—let us say white balls represent the reference allele and black balls the alternative allele. Whenever a ball is drawn, it is replaced to the urn and another $c$ balls of the same *c*olor are added, analogous to the polymerase copying an allele.

As we have two strands each for both alleles, a natural choice of starting alleles would be $w_t = b_t = 2$. We here subscript the values with $_t$ to denote the theoretical nature of the parameter choice, as opposed to practical values of $w_p, b_p$ introduced below. And as one strand is added with every copy operation, we set $c = 1$ (Figure S 1A). This gives us a slight peak around the alternative allele frequency of 0.5 (Figure S 1A), which we can also make out in the real data (Figure 1B of the main manuscript).

However, it does not capture the allele dropout peaks at the allele frequencies of 0 and 1 seen there. A likely reason for these dropout peaks is, that not every position on every allele is equally accessible to amplification initiation [Picher et al., 2016]. I.e. mechanistically, some close-by site to the one under consideration must be single-stranded at some point during the continuous amplification reaction (at a constant temperature), to be accessible for an initial priming that will start the amplification of the respective allele for that region. To account for this lower initiation accessibility, we can introduce a scaling factor $f$ for the starting frequencies of the two alleles, with $0 <= f <= 1$,

[6] The notation used here is analogous to Equation 6.28 in Section 6.2.4 of Johnson et al. [2005]. Also see Equation S 7 in the above footnote for its relation to $\alpha_1, \alpha_2$ in Lodato et al. [2015].

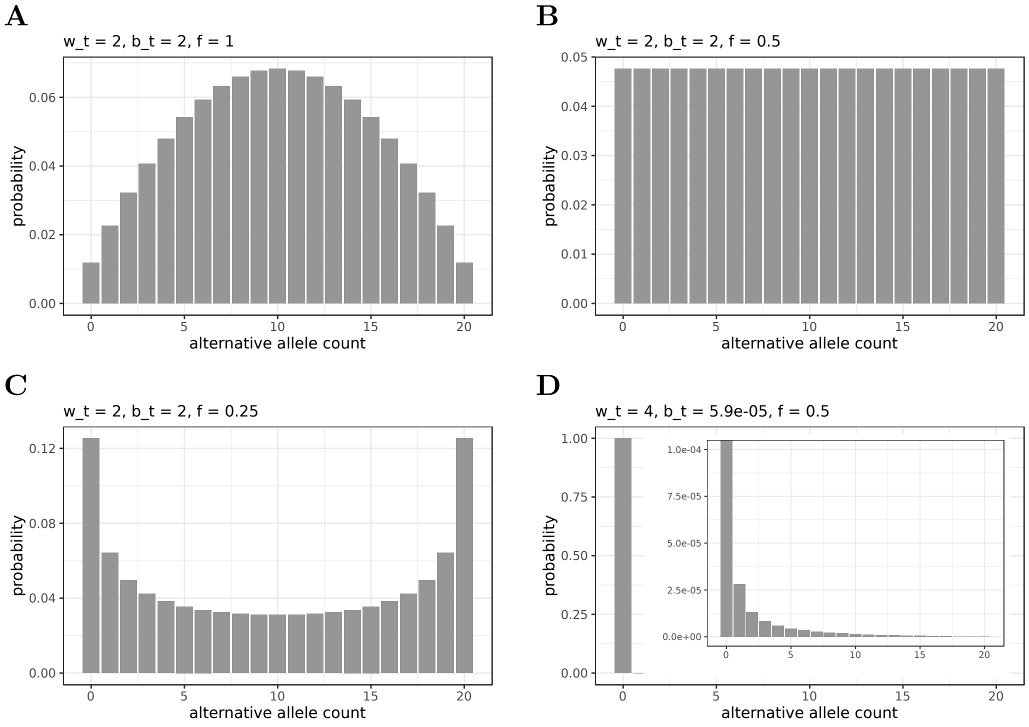

Figure S 1: Beta-binomial distributions with different shape parameters, that correspond to our theoretical numbers of starting allele strands ($w_t = b_t = 2$) scaled to different amplification accessibility by $f$. **A** Perfect accessibility of both strands of both alleles for amplification ($f = 1$) and the resulting alternative allele frequencies. **B** Half that accessibility ($f = 0.5$), resulting in the uniform distribution. **C** For any accessibility scaling value below $\frac{1}{2}$ (e.g. $f = 0.25$), the beta-binomial distribution will give maxima at alternative allele frequencies 0 and 1. **D** The heavily non-symmetrical case ($w_t \gg b_t$), analogous to homozgyous reference genotypes. Instead of the distribution currently used in ProSolo, which was fit to data and compounds sequencing errors with amplification errors (Figure 1A in the main manuscript), this distribution makes the following theoretical assumptions: It sets $w_t = 4$ for the 4 reference strands available for amplification and $b_t$ to four times the known $\Phi 29$ error rate for drawing / introducing an error alternative allele instead of the reference allele. The inset zooms into the y-axis to show details of the tail of the distribution.

to scale down the initially available allele count: if strands are already separated for this allele and amplification initiation works perfectly for both strands ($f = 1$), then we would get $w_p = w_t * f = 2 * 1 = 2$; whereas at the other extreme of $f = 0$, amplification initialisation of the allele would be impossible. Then, for $f = \frac{1}{2}$, we get the uniform distribution ($w_p = w_t * f = \frac{1}{2} * 2 = 1$, Figure S 1B), and for all $f < \frac{1}{2}$, we will get distributions with peaks at 0 and 1 (e.g. Figure S 1C).

If we then consider a mixture of sites with different $f$, we would expect distributions as observed e.g. in Figure 1B of the main manuscript; and a reasonable approximation of this is the mixture of the two symmetrical beta-binomials as fitted by Lodato et al. [2015], one with $f \approx 0.40 < \frac{1}{2}$ and one with $f \approx 0.67 > \frac{1}{2}$ (i.e. the intersects of $\alpha_2$ and $\alpha_1$ plus their slight slope-adjustments, as given in Table S 1).

##### S 1.2.2 Outlook: more realistic beta-binomial distributions

However, something that this does not capture is possible asymmetry of the distributions, i.e. the case of $w \neq b$. While the theoretical starting amounts of strands of both alleles in the heterozygous case should remain the same ($w_t = b_t = 2$), we would have to introduce allele-specific scaling factors $f_w \neq f_b$ that capture the different availability of the two alleles for amplification (e.g. because one of the alleles is more densely wrapped around Histones than the other). As a result we would get $w_p = w_t * f_w \neq f_b * b_t = b_p$, and instead of the one shape parameter in the symmetrical case, we would then have to fit the distributions with a second shape parameter. I.e. instead of Equation S 7, we would get:

$$\begin{aligned} \alpha_1 &= w_p = w_t * f_w = 2f_w \\ \beta_1 &= b_p = b_t * f_b = 2f_b \end{aligned} \tag{S 9}$$

However, assuming that the dropout of both alleles—reference and alternative—is equally likely, this asymmetry will average out over all heterozygous sites, and the symmetry assumption remains a justified reduction in parameters to fit when fitting a mixture of two beta-binomials.

This is somewhat different for the homozygous case. Here, Lodato et al. [2015] already modelled the allele frequency deviations ($> 0$ for homozygous reference) as an asymmetric beta-binomial distribution. This gives a reasonable fit to the data and makes sense to use, as we would expect these deviations to be the result of amplified polymerase errors from the MDA. While we just use the distributions fit by Lodato et al. [2015] in our initial implementation we present here, this could in the future be replaced by a theoretically derived beta-binomial distribution. For example, for the homozygous reference case, a very low $b_t$ (the probability of "choosing" a previously non-existing error allele) could be determined by (four times) the known $\Phi29$ polymerase error rate, whereas $w_t$ would be set to 4 to account for four reference strands available for copying (Figure S 1D). As $f$ would in this case affect both the reference ($w$) and the error allele ($b$) alike, it could be set to some identical value for both, e.g. to an average accessibility of 0.5 (Figure S 1D). This would avoid compounding the uncertainty due to MDA with sequencing errors, that we already account for separately in our latent variable model through base calling quality scores and is one very likely source of the current mis-classifications of ground truth heterozygous sites as homozygous (Figure S 9).

##### S 1.3 Reads: Sampling Alternative Nucleotide Frequencies in the Sequencing Process

Let $\mathfrak{R}$ ($=\mathfrak{R}(g)$) denote all reads covering a site $g$, with $R_i \in \mathfrak{R}$ denoting an individual read out of the $l$ reads aligning to that site ($l \in \mathbb{N}, i \in \{1, 2, \ldots, l\}$). Formally, we index positions in aligned reads by the respective genome indices $g$, but we omit the index $g$

for readability. Working with a given $\rho$, one can assume independence among the reads in the respective sample (while, of course, given $\theta$, one can assume independence only in bulk, due to the necessary amplification and the resulting coverage nonuniformity in single cell samples). That is

$$L(\rho \mid \mathfrak{R}) \propto P(\mathfrak{R} \mid \rho) = \prod_{R \in \mathfrak{R}} P(R \mid \rho) \propto \prod_{R \in \mathfrak{R}} L(\rho \mid R) \tag{S 10}$$

both for bulk and single cell sequencing. Let $S_{R_i}[g]$ be the nucleotide determined by the sequencer for position $g$ of read $R_i$ and $Q_{R_i}[g]$ be the Phred quality score for the nucleotide at that position of read $R_i$[7]. Let further $M_{R_i}[g]$ be the Phred alignment quality score[8] provided by the aligner. We now define $Z_i[g]$ ($i \in \{1, \dots, l\}$) to be the collection of the observable variables for the alignment of one of $l$ reads $R_i \in \mathfrak{R}$ to a genome position $g$. Omitting index $g$, this is the vector $Z_i = \langle S_{R_i}, Q_{R_i}, M_{R_i} \rangle$ and drawing on this information, we now introduce hyperparameters analogously to the model for tumor-normal variant calling as outlined by Köster et al. [2019] for Varlociraptor. These will be the latent variables $\omega_i \in \{0, 1\}$ to model correct placement of an alignment ($\omega_i = 1$ if correct and $\omega_i = 0$ otherwise) and $\xi_i \in \{0, 1\}$ to specify whether an alignment $Z_i$ is associated with the alternative nucleotide ($\xi_i = 1$) or not ($\xi_i = 0$). Taken together, they give three cases

$$(\omega_i, \xi_i) = \begin{cases} (1,1) : R_i \text{ mapped correctly, alternative nucleotide} \\ (1,0) : R_i \text{ mapped correctly, reference nucleotide} \\ (0,\{0,1\}) : R_i \text{ not mapped correctly, nucleotide irrelevant} \end{cases} \tag{S 11}$$

[7] As above, we now omit $g$ for $S_{R_i}$ and $Q_{R_i}$ for ease of notation and reading, as we will for $M_{R_i}$. Also note that the value of $Q_{R_i}[g]$ can be adjusted by a base quality score recalibration that accounts for systematic base calling errors, before using it as input to ProSolo.

[8] Also called mapping quality; represented by the MAPQ tag in the sequence alignment (SAM) format specifications v1 (2a802cd) and by the MQ tag in the variant call format (VCF) specification v4.3 (2a802cd). As before, we will omit the position $g$ from notation from this point on.

We thus model the two major sources of uncertainty on the level of the individual alignment $Z_i$, 1) *alignment uncertainty* and 2) *typing uncertainty*, by associating every alignment $Z_i$ with these two binary-valued, latent variables, reflecting uncertainty hyperparameters, $\omega_i$ and $\xi_i$:

*First*,

$$\omega_i \sim \text{Bernoulli}\,(\pi_i) \;\text{ for } i = 1, \ldots, l \tag{S 12}$$

where $\pi_i$ is the (Phred scaled) posterior probability proportional to $M_{R_i}$ provided by the aligner. We can compute it with

$$\pi_i = 10^{-M_{R_i}/10} \tag{S 13}$$

*Second*,

$$\xi_i \sim \text{Bernoulli}\,(\rho) \;\text{ for } i = 1, \ldots, l \tag{S 14}$$

which reflects that sampling a fragment from the locus that bears the alternative nucleotide agrees with the probability to sample a genome copy with the alternative nucleotide from the pool of amplified DNA.

Whether $\xi_i$ is 1 or 0 is generally not evident from the observed $Z_i$, due to typing uncertainty. We formally define[9]

$$Z_i \mid \omega_i = 1, \xi_i = 0 \sim a(Z_i) \quad \text{ and } \quad Z_i \mid \omega_i = 1, \xi_i = 1 \sim p(Z_i) \tag{S 15}$$

[9] If $\omega_i = 0$, that is the alignment is incorrect, $Z_i \mid \omega_i = 0 \sim 1$ reflects that $Z_i$ has no influence on the posterior probability distribution of $\rho$.

where $a, p$ are probability distributions over correct $S_{R_i}$ when the alternative nucleotide is either $a$bsent or $p$resent in read $R_i$ and $a(\cdot)$ and $p(\cdot)$ are the respective functions to sample from this distribution for a particular read. With these probability distributions, we can account for any number of factors for which we know how they affect these probability distributions—but if we lack any further information for an alternative nucleotide, we can assume $Q_{R_i}$ to be a specification of these distributions for $R_i$, accounting for the uncertainties of the sequencing process. With $x \in \{A, C, G, T\}$ the alternative nucleotide under inspection, we can thus compute:

$$a(Z_i) = \begin{cases} 10^{-Q_{R_i}/10} & \text{if } S_{R_i} = x \\ 1 - 10^{-Q_{R_i}/10} & \text{if } S_{R_i} \neq x \end{cases} \tag{S 16}$$

$$p(Z_i) = 1 - a(R_i) \tag{S 17}$$

##### S 1.4 Single cell latent variable model

By applying the Chapman-Kolmogorov equation with our binary hyperparameters to the single cell case[10], we get

$$L(\rho_s \mid Z_i^s) \propto \; P(Z_i^s \mid \rho_s) = \int_{\omega_i^s, \xi_i^s} P(Z_i^s \mid \omega_i^s, \xi_i^s) \times P(\omega_i^s, \xi_i^s \mid \rho_s) \; d(\omega_i^s, \xi_i^s) \tag{S 18}$$

which fans out to the following cases from equation S 11:

[10] We will only be looking at single cell samples in the following, as the case of $\rho_b = \theta_b$ is the simpler one and analogous without the whole genome amplification considerations. Please also refer to the tumor and normal sample derivations in Köster et al. [2019], but note that we need no purity parameter, here.

$$
\begin{aligned}
P(Z_i^s \mid \rho_s) = P(Z_i^s \mid \omega_i^s = 0) \times P(\omega_i^s = 0 \mid \rho_s) \\
+ P(Z_i^s \mid \omega_i^s = 1, \xi_i^s = 0) \times P(\omega_i^s = 1, \xi_i^s = 0 \mid \rho_s) \\
+ P(Z_i^s \mid \omega_i^s = 1, \xi_i^s = 1) \times P(\omega_i^s = 1, \xi_i^s = 1 \mid \rho_s)
\end{aligned}
\tag{S 19}
$$

Knowing that the mapping probability of reads is independent from the given (distorted) allele frequency $\rho_s$, and assuming a sampling probability of 1 for any mismapped read, this becomes:

$$
\begin{aligned}
P(Z_i^s \mid \rho_s) = P(\omega_i^s = 0) \\
+ P(Z_i^s \mid \omega_i^s = 1, \xi_i^s = 0) \times P(\omega_i^s = 1) \times P(\xi_i^s = 0 \mid \rho_s) \\
+ P(Z_i^s \mid \omega_i^s = 1, \xi_i^s = 1) \times P(\omega_i^s = 1) \times P(\xi_i^s = 1 \mid \rho_s)
\end{aligned}
\tag{S 20}
$$

With Equation S 12 for $\omega_i^s$, Equation S 14 for $\xi_i^s$ and Equations S 16 and S 17 for the conditional probabilities of the data given an absence or presence of the observed nucleotide, we get:

$$
\begin{aligned}
P(Z_i^s \mid \rho_s) = (1 - \pi_i^s) \\
+ \ a^s(Z_i^s) \times \pi_i^s \times (1 - \rho_s) \\
+ \ p^s(Z_i^s) \times \pi_i^s \times \rho_s
\end{aligned}
\tag{S 21}
$$

As the $l$ reads are conditionally independent, given a particular (distorted) alternative nucleotide frequency $\rho_s$ (also see Equation S 10), we can calculate the likelihood of such a particular $\rho_s$ with:

$$
L(\rho_s \mid \boldsymbol{Z}^s) \propto P(\boldsymbol{Z}^s \mid \rho_s) = \prod_{i=1}^{l} P(Z_i^s \mid \rho_s) \tag{S 22}
$$

With the resulting likelihoods for $\rho_s \in [0, 1]$, we can calculate the likelihood of a particular $\theta_s$:

$$L(\theta_s \mid \boldsymbol{Z}^s) \propto P(\boldsymbol{Z}^s \mid \theta_s) = \sum_{k=0}^{l} \left\{ P(\boldsymbol{Z}^s \mid \rho_s = \frac{k}{l}) \times P(\rho_s = \frac{k}{l} \mid \theta_s) \right\} \tag{S 23}$$

For $\theta_s \in \{0, \frac{1}{2}, 1\}$, with equation S 5 specifying the cases of $\theta_s$ and Equation 2 further specifying $P(\rho_s \mid \theta_s)$ for MDA, this gives us:

$$\begin{aligned}
P(\boldsymbol{Z}^s \mid \theta_s = 0) &= \sum_{k=0}^{l} \left\{ P(\boldsymbol{Z}^s \mid \rho_s = \frac{k}{l}) \times \mathfrak{BB}_{l,\alpha,\beta}(\rho_s = \frac{k}{l}) \right\} \\
P(\boldsymbol{Z}^s \mid \theta_s = \frac{1}{2}) &= \sum_{k=0}^{l} \left\{ P(\boldsymbol{Z}^s \mid \rho_s = \frac{k}{l}) \times \left[ w(l) \times \mathfrak{BB}_{l,\alpha_1,\alpha_1}(\rho_s = \frac{k}{l}) + \right.\right. \\
&\qquad \left.\left. (1 - w(l)) \times \mathfrak{BB}_{l,\alpha_2,\alpha_2}(\rho_s = \frac{k}{l}) \right] \right\} \\
P(\boldsymbol{Z}^s \mid \theta_s = 1) &= \sum_{k=0}^{l} \left\{ P(\boldsymbol{Z}^s \mid \rho_s = \frac{k}{l}) \times \mathfrak{BB}_{l,\beta,\alpha}(\rho_s = \frac{k}{l}) \right\}
\end{aligned} \tag{S 24}$$

For the current implementation, we use the empirical values for $\alpha(l), \beta(l), w(l), \alpha_1(l), \alpha_2(l)$ as deduced by Lodato et al. [2015] (Table S 1). But as mentioned above, these could be learned from a dataset at hand whenever high-confidence sets of heterozygous and homozygous sites are available (e.g. by calling variants on a background bulk and cross-referencing with dbSNP).

#### S 1.5 Bulk background: Accounting for Amplification Errors and Allele Dropout

Using the above model for a genome position $g$, we can compute the likelihood of any combination of possible underlying alternative nucleotide frequencies in the single cell

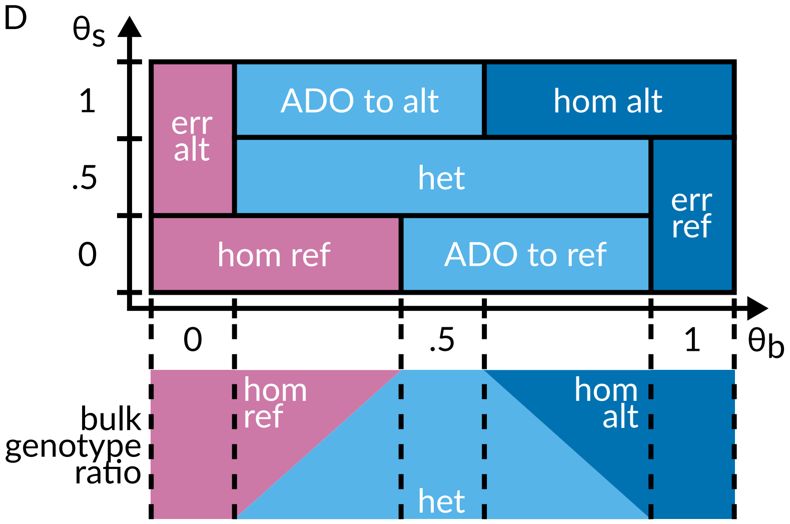

Figure S 2: It depicts single cell events at a particular genomic site, defined by combinations of the possible true alternative nucleotide frequencies in single cell sequencing ($\theta_s$) and bulk sequencing samples ($\theta_b$) in our probabilistic model. The bulk is always assumed to be a combination of a maximum of two genotypes at a particular site, generating all possible $\theta_b$ (bottom panel). A tabular representation of the same event definitions is given in Table S 2 below (this is a larger copy of Figure 1D in the main manuscript, reproduced here for easier cross-reference with that table).
ADO – allele dropout, alt – alternative, err – error, het – heterozygous, hom – homozygous, ref – reference

Table S 2: Single cell events at a particular genomic site are defined by combinations of the possible true alternative nucleotide frequencies in single cell sequencing ($\theta_s$) and bulk sequencing samples ($\theta_b$) in our probabilistic model. A graphical representation of the same event definitions is given in Figure S 2.

| single cell event | $\theta_s(x)$ | $\theta_b(x)$ | implicit single cell genotype |
|---|---|---|---|
| $\boldsymbol{E}_{\text{hom ref}}$ | 0 | $[0, \frac{1}{2})$ | homozygous ref |
| $\boldsymbol{E}_{\text{ADO to ref}}$ | 0 | $[\frac{1}{2}, 1)$ | heterozygous |
| $\boldsymbol{E}_{\text{err ref}}$ | $\{0, \frac{1}{2}\}$ | 1 | homozygous alt |
| $\boldsymbol{E}_{\text{err alt}}$ | $\{\frac{1}{2}, 1\}$ | 0 | homozygous ref |
| $\boldsymbol{E}_{\text{het}}$ | $\frac{1}{2}$ | $(0, 1)$ | heterozygous |
| $\boldsymbol{E}_{\text{ADO to alt}}$ | 1 | $(0, \frac{1}{2}]$ | heterozygous |
| $\boldsymbol{E}_{\text{hom alt}}$ | 1 | $(\frac{1}{2}, 1]$ | homozygous alt |

ADO – allele dropout, alt – alternative, err – error, het – heterozygous, hom – homozygous, ref – reference and the bulk sequencing sample, given the respective data. All the possible combinations of allele frequencies in the two samples define a two-dimensional space of allele frequencies $\boldsymbol{E}$ (Equation 4, Figure S 2, Table S 2). We then define mutually exclusive subspaces of $\boldsymbol{E}$ as single cell events, for example the dropout of an alternative allele across allele frequencies $\boldsymbol{E}_{\text{ADO to ref}} = \{0\}_{\theta_s} \times [\frac{1}{2}, 1)_{\theta_b}$, providing a set of events $\mathfrak{E}$. Assuming that there is no informative prior information about different allele frequencies (i.e. assuming a flat prior across all possible allele frequencies in both samples), we can then calculate the likelihood of such an event as the product of the likelihoods of the allele frequency ranges in the two samples. For $\boldsymbol{E}_{\text{ADO to ref}}$—and using Equation S 24 to calculate the likelihoods of the allele frequency ranges—this for example becomes:

$$\begin{aligned} P(\boldsymbol{Z}^s, \boldsymbol{Z}^b \mid \boldsymbol{E}_{\text{ADO to ref}}) &= P(\boldsymbol{Z}^s \mid \theta_s = 0) \times P(\boldsymbol{Z}^b \mid \theta_b \in [\frac{1}{2}, 1)) \\ &= \sum_{k=0}^{l} \left\{ P(\boldsymbol{Z}^s \mid \rho_s = \frac{k}{l}) \times \mathfrak{B}\mathfrak{B}_{l,\alpha,\beta}(\rho_s = \frac{k}{l}) \right\} \times \\ &\quad \sum_{m=\frac{n}{2}}^{n-1} P(\boldsymbol{Z}^b \mid \theta_b = \frac{m}{n}) \end{aligned} \tag{S 25}$$

Here, we use the fact that for the bulk sample without amplification, we can assume that $\theta_b = \rho_b$ and thus use a simpler form of Equation S 23. Also, we take $n$ to be the total number of bulk reads and $m$ to be the possible numbers of reads with the alternative allele for this event—analogous to $k$ and $l$ for the single cell sample, and amounting to only taking point estimates of the likelihoods at possible alternative allele counts given the total coverage of the examined genomic site.

As all the defined events (Equation 5) are mutually exclusive (Figure S 2, Table S 2), the sum of the likelihoods of all events yields the marginal probability:

$$
\begin{aligned}
P(\boldsymbol{Z}^s, \boldsymbol{Z}^b) = P(\boldsymbol{Z}^s, \boldsymbol{Z}^b \mid \boldsymbol{E}) = & \sum_{\boldsymbol{E}_e \in \mathfrak{E}} P(\boldsymbol{Z}^s, \boldsymbol{Z}^b \mid \boldsymbol{E}_e) \\
= & \, P(\boldsymbol{Z}^s, \boldsymbol{Z}^b \mid \boldsymbol{E}_{\text{hom ref}}) \; + \; P(\boldsymbol{Z}^s, \boldsymbol{Z}^b \mid \boldsymbol{E}_{\text{hom alt}}) + \\
& P(\boldsymbol{Z}^s, \boldsymbol{Z}^b \mid \boldsymbol{E}_{\text{het}}) + \\
& P(\boldsymbol{Z}^s, \boldsymbol{Z}^b \mid \boldsymbol{E}_{\text{err alt}}) \; + \; P(\boldsymbol{Z}^s, \boldsymbol{Z}^b \mid \boldsymbol{E}_{\text{ADO to alt}}) + \\
& P(\boldsymbol{Z}^s, \boldsymbol{Z}^b \mid \boldsymbol{E}_{\text{err ref}}) \; + \; P(\boldsymbol{Z}^s, \boldsymbol{Z}^b \mid \boldsymbol{E}_{\text{ADO to ref}})
\end{aligned}
\tag{S 26}
$$

Then, using Equation S 22 for the likelihood of $\rho_s$ in the single cell sample and the equivalent relationship for the likelihood of $\theta_b$ in the bulk sample (with $j$ as an index running through all bulk sample reads), we get:

$$
\begin{aligned}
P(\boldsymbol{Z}^s, \boldsymbol{Z}^b \mid \boldsymbol{E}_{\text{ADO to ref}}) = & \sum_{k=0}^{l} \left\{ \prod_{i=1}^{l} P(Z_i^s \mid \rho_s = \frac{k}{l}) \times \mathfrak{BB}_{l,\alpha,\beta}(\rho_s = \frac{k}{l}) \right\} \times \\
& \sum_{m=\frac{n}{2}}^{n-1} \prod_{j=1}^{n} P(Z_j^b \mid \theta_b = \frac{m}{n})
\end{aligned}
\tag{S 27}
$$

Using Equation S 21 for the likelihood of an individual read in the single cell sample given a $\rho_s$ and the respective formulation for an individual read in the bulk sample given a $\theta_b$, this will calculate the likelihood of the `ADO to ref` event as:

$$
\begin{aligned}
& P(\boldsymbol{Z}^s, \boldsymbol{Z}^b \mid \boldsymbol{E}_{\text{ADO to ref}}) = \\
& \quad \sum_{k=0}^{l} \left\{ \prod_{i=1}^{l} \left( (1 - \pi_i^s) + a^s(Z_i^s) \times \pi_i^s \times (1 - \frac{k}{l}) + p^s(Z_i^s) \times \pi_i^s \times \frac{k}{l} \right) \times \right. \\
& \qquad \left. \mathfrak{BB}_{l,\alpha,\beta}(\rho_s = \frac{k}{l}) \right\} \times \\
& \quad \sum_{m=\frac{n}{2}}^{n-1} \left\{ \prod_{j=1}^{n} \left( (1 - \pi_j^b) + a^b(Z_j^b) \times \pi_j^b \times (1 - \frac{m}{n}) + p^b(Z_j^b) \times \pi_j^b \times \frac{m}{n} \right) \right\}
\end{aligned}
\tag{S 28}
$$

Analogously, the likelihoods for all of the events as specified in Figure S 2 can be computed and their sum according to Equation S 26 is the marginal probability. We can use the marginal probability to calculate the posterior probability of the `ADO to ref` event with:

$$
\begin{aligned}
&P(\boldsymbol{E}_{\text{ADO to ref}} \mid \boldsymbol{Z}^s, \boldsymbol{Z}^b) = \\
&\quad \frac{1}{\sum_{\boldsymbol{E}_e \in \mathfrak{E}} P(\boldsymbol{Z}^s, \boldsymbol{Z}^b \mid \boldsymbol{E}_e)} \times \\
&\quad \sum_{k=0}^{l} \left\{ \prod_{i=1}^{l} \left( (1-\pi_i^s) + a^s(Z_i^s) \times \pi_i^s \times (1-\frac{k}{l}) + p^s(Z_i^s) \times \pi_i^s \times \frac{k}{l} \right) \times \right. \\
&\quad\quad \left. \mathfrak{B}\mathfrak{B}_{l,\alpha,\beta}(\rho_s = \frac{k}{l}) \right\} \times \\
&\quad \sum_{m=\frac{n}{2}}^{n-1} \left\{ \prod_{j=1}^{n} \left( (1-\pi_j^b) + a^b(Z_j^b) \times \pi_j^b \times (1-\frac{m}{n}) + p^b(Z_j^b) \times \pi_j^b \times \frac{m}{n} \right) \right\}
\end{aligned}
\tag{S 29}
$$

With analogous equations, we can compute the posterior probability for all of the events defined by ProSolo (Figure S 2, Table S 2). Each of them is reported in the output VCF or BCF file. Based on the implicit single cell genotypes as given in Table S 2, we can then compute posterior probabilities for genotypes in the single cell by summing the posterior probabilities of all the events that imply each genotype. For the heterozygous case (light blue fields in Figure S 2), this would be:

$$
\begin{aligned}
P(\text{heterozygous}_s \mid \boldsymbol{Z}^s, \boldsymbol{Z}^b) = &\, P(\boldsymbol{E}_{\text{ADO to ref}} \mid \boldsymbol{Z}^s, \boldsymbol{Z}^b) + \\
&\, P(\boldsymbol{E}_{\text{het}} \mid \boldsymbol{Z}^s, \boldsymbol{Z}^b) + \\
&\, P(\boldsymbol{E}_{\text{ADO to alt}} \mid \boldsymbol{Z}^s, \boldsymbol{Z}^b)
\end{aligned}
\tag{S 30}
$$

The genotype with the maximum probability among the three possible genotypes is then chosen. Similarly, we can also compute posterior probabilities for any compound event. For example, the overall posterior probability of an allele dropout at a particular site (`ADO` events in Figure S 2 and Table S 2) is:

$$P(\text{allele dropout} \mid \boldsymbol{Z}^s, \boldsymbol{Z}^b) = P(\boldsymbol{E}_{\text{ADO to ref}} \mid \boldsymbol{Z}^s, \boldsymbol{Z}^b) + P(\boldsymbol{E}_{\text{ADO to alt}} \mid \boldsymbol{Z}^s, \boldsymbol{Z}^b) \tag{S 31}$$

Another example is the probability of the presence of any alternative allele at a site (all blue fields in Figure S 2, Equation 10 in the main manuscript). This posterior probability is the one used for the main benchmarking of ProSolo against the other existing single cell specific variant callers (MonoVar, SCAN-SNV, SCcaller, and SCIPhI), as SCcaller and SCIPhI only call the presence (vs. absence) of an alternative nucleotide and this is thus the only measure we can compare all tools on.

#### S 1.6 Implementation

ProSolo is an easy-to-use command-line tool (following usability standards [Taschuk and Wilson, 2017]) and its source code is available at `https://github.com/prosolo/prosolo`, including instructions for an easy installation (via Bioconda [Grüning et al., 2018]). The main contribution in terms of software that ProSolo's development provided, is the implementation of its comprehensive statistical model that jointly looks at a single-cell and a bulk sample. This was implemented as a part of Varlociraptor, a variant calling library implemented in Rust [Köster et al., 2019]. Varlociraptor already makes use of all available sources of information regarding biases of normal sequencing data, induced by ambiguous alignments and sequencing errors, but had to be extended to accommodate this type of sample combination and to allow for accounting for amplification bias. Because Varlociraptor had originally been specializing in indels and structural variants, implementing ProSolo also meant further investment in expanding Varlociraptor's capabilities for basic SNV analysis and required some improvements in the handling of sequencing alignment data in the upstream library rust-htslib [rust htslib], a Rust wrapper around htslib [htslib]. Further, we optimized Varlociraptor's calculations wherever possible, for example by reducing the number of redundant likelihood calculations and by caching frequently reused values. And finally, numerical issues required an addition to the upstream library rust-bio [Köster, 2016].

**False Discovery Rate (FDR) Control of Event Probabilities** Last—but not least—ProSolo can control the false discovery rate of any set of single cell events that a user defines. At any scale of high throughput sequencing data (from targeted panels over whole exome sequencing to whole genome sequencing), we generate a large amount of event calls and thus have to control for false discoveries, e.g. by controlling the false discovery rate [Benjamini and Hochberg, 1995]. Using Varlociraptor's functionality [Köster et al., 2019], we can estimate thresholds on our posterior probabilities based on the approach described by Müller et al. [2004, 2006]. For ProSolo, we extended Varlociraptor's capabilities from estimating such thresholds for the posterior probability of a single event to estimating thresholds for any combination of the above defined events. For example, to control the false discovery rate for allele dropout events, we jointly obtain a threshold for the combination of the two ADO events in Figure 1D. Or, to control the false discovery rate in identifying the presence of an alternative allele, we jointly obtain a threshold for the combination of all the respective events in Equation 10. This functionality, like the improvements mentioned in Section S 1.6, is now also available in Varlociraptor's subcommands.

### S 2 Benchmarking

All code used for benchmarking is available at: `https://www.github.com/prosolo/benchmarking_prosolo`.

##### S 2.1 Datasets and Ground Truths

We have performed our benchmarking analyses on two datasets, one previously published for the evaluation of SCcaller by Dong et al. [2017] and one generated especially for this Benchmarking (see Figure 2 in the main manuscript). For them, we generated different kinds of ground truths:

**Whole genome sequencing of almost identical kin-cells from a cell-line [Dong et al., 2017].** The first dataset is from the publication of a previous variant caller for single cell sequencing data generated using MDA, SCcaller (Dong et al. [2017]; Figure 2A). They started with one initial cell from a cell line and grew it in several steps (a complete overview of their setup is given in Figure 1 of Dong et al. [2017]). After an initial mini-expansion, they selected one single cell as the founder for their secondary IL expansion into 20-30 cells. From this they extracted two single cells (IL-11 and IL-12) that were sequenced after multiple displacement amplification (MDA). Both cells have a minimum coverage of five of at least 80% of genomic sites (with only about 90% of the used reference genome represented, as this includes a largely uncovered chromosome Y and alternative haplotype contigs which would only be covered if the particular haplotype is present), with IL-11 having a more even coverage that leads to more sites with a usable coverage (Figure S 3A). The rest of the kindred cells from that IL clone was then used as a bulk sequencing sample without MDA (IL-1C), showing a coverage profile similar to the whole genome amplified single cells (Figure S 3A). We generated the ground truth genotypes for this dataset from the kindred clone IL-1C, which should be only very few cell divisions away from the single cells, and thus have almost no difference in the somatic mutations acquired. The ground truth genotype was generated separately for variant sites and homozygous reference sites: for variant sites, GATK HaplotypeCaller was run, followed by variant quality score recalibration; for homozygous reference sites, we used bcftools mpileup to identify sites with a coverage above 25 but no alternative nucleotide present. This bulk sample (IL-1C) was only used as a ground truth and was not provided as input to any of the software compared here. Three more distant clones (C1, C2, C3 in Figure 2A), grown from the cells after the initial mini-expansion, were jointly used as a further bulk sample for SCcaller and ProSolo (see Software and Parameters below). With three separate bulk sequencing runs, their joint coverage of about 50 for 80% of genomic sites is much higher than for the MDA single cells and the unamplified IL-1C bulk (Clones in Figure S 3A).

**Whole exome sequencing of five human granulocytes with a pedigree ground truth.** For the second benchmarking dataset, Blood was taken from a patient after informed consent and granulocytes were sorted out in several steps: Initially, CD3- cells were captured via Magnetic-Activated Cell Sorting (MACS); among them, CD66b+ granulocytes were filtered out via Fluorescence Activated Cell Sorting (FACS) (Figure 2B). Single cells from that cell population were isolated using a Fluidigm C1 microfluidics chip and subjected to multiple displacement amplification (MDA) using the GE Healthcare GenomiPhi Kit in its version adapted to that microfluidic chip. With a panel of 16 custom primer pairs covering different genes across chromosomes used in a quantitative real-time PCR, we selected granulocytes where at least 15 of these loci were properly amplified. For those cells, we performed whole exome capture using the Agilent SureSelectXT Human All Exon V5+UTRs Kit. The paired-end sequencing library (2x101bp) was then prepared with the TruSeq V3 Kit and sequenced on a HiSeq 2500. From the remaining sorted cell population, we extracted bulk DNA and submitted it to the same whole exome capture and DNA sequencing without MDA.

For the ground truth of this dataset, we could leverage previously published bulk whole exome sequencing data from the same person, their parents and three siblings (Hoell et al. [2014]; Figure 2B). For ground truth alternative allele calls, we ran three

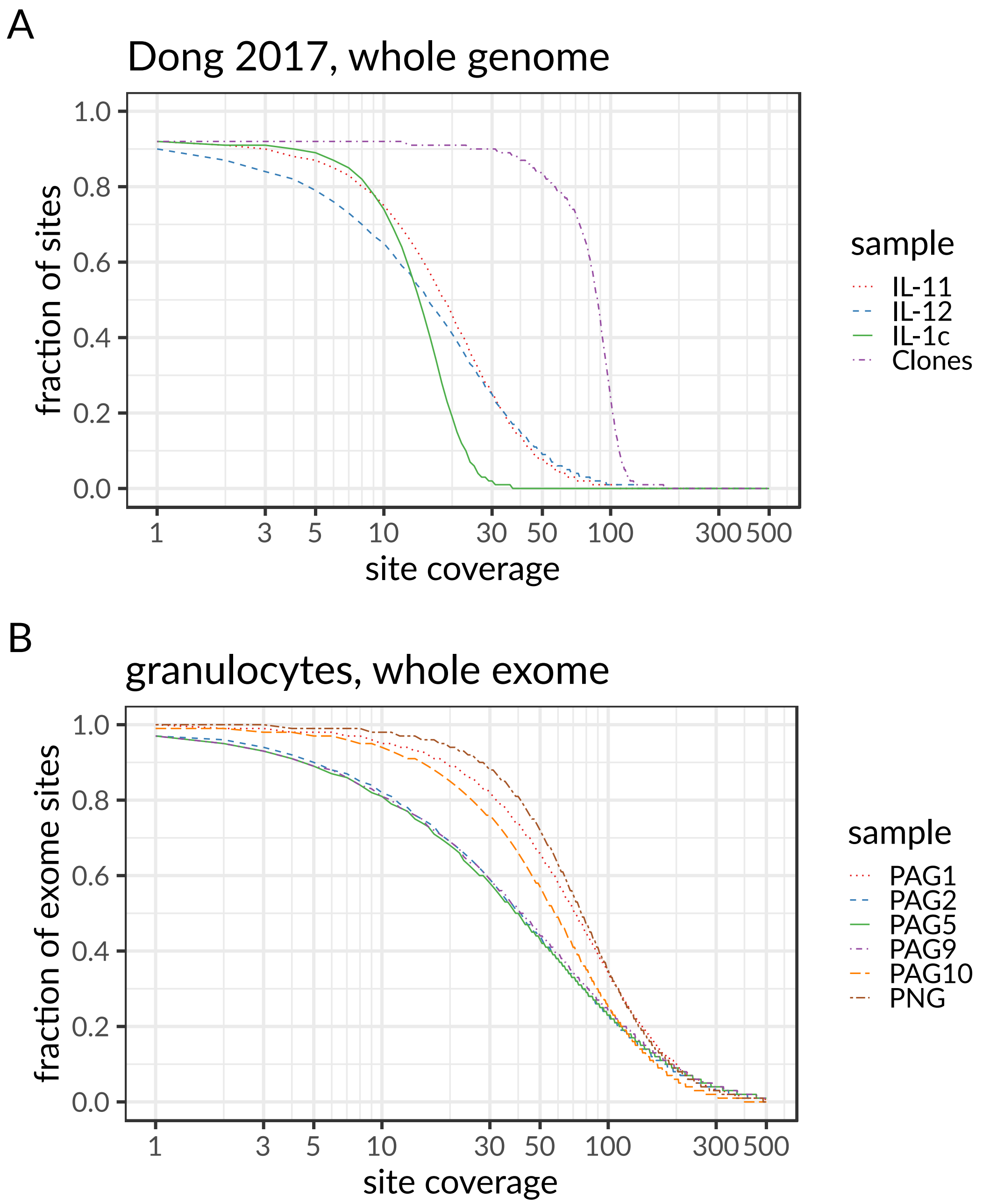

Figure S 3: Sequencing coverage of targeted genomic sites for multiple displacement amplified (MDA) single cells and respective bulk samples. Panel **A** shows the coverage of the single cells (IL-11, IL-12), the kindred ground truth clone (IL-1c) and the more distant background bulk (Clones C1, C2, and C3 joined into one sample) of the dataset from Dong et al. [2017]. Panel **B** shows the coverage of the single cells (PAG1, PAG2, PAG5, PAG9, PAG10) and the background bulk (PNG) of the granulocyte data generated for this evaluation.

pedigree-aware variant callers on the family whole exome data. To be able to run them reproducibly, we made all three of them available via bioconda [Grüning et al., 2018]:

- BEAGLE 4.0 [Browning and Browning, 2013], bioconda version: `beagle=4.0_06Jun17=1`
- polymutt [Li et al., 2012], bioconda version: `polymutt=0.18=0`
- FamSeq in MCMC (`-method 3`) mode [Peng et al., 2013, 2014]), bioconda version: `famseq=1.0.3=0`

From their calls, we created a consensus by only including sites where either all callers agree on a genotype or where a maximum of one caller has a *missing* (as opposed to different genotype) call, while the other two callers agree on the genotype (Figure S 4). The use of the full pedigree ensures high confidence genotypes—however, please bear in mind that these represent the germline genotype. Single cells will have accumulated somatic genotypes that differ from the germline and we are bound to mis-interpret those as false positive calls in our ground truth comparison. At the same time, the number of such somatic variants will be small compared to the number of germline variants, and should hence keep our overall ground truth comparison valid.

##### S 2.2 Software and Parameters

ProSolo was compared against the available tools for single nucleotide variant calling in single cell sequencing data from multiple displacement amplified (MDA) DNA, MonoVar [Zafar et al., 2016], SCAN-SNV [Luquette et al., 2019], SCcaller [Dong et al., 2017], and SCIPhI [Singer et al., 2018].

For ProSolo (version `0.6.1` installed via bioconda), each single cell was called against the respective cell's bulk (the joined clones 1, 2 and 3 for the [Dong et al., 2017] dataset,

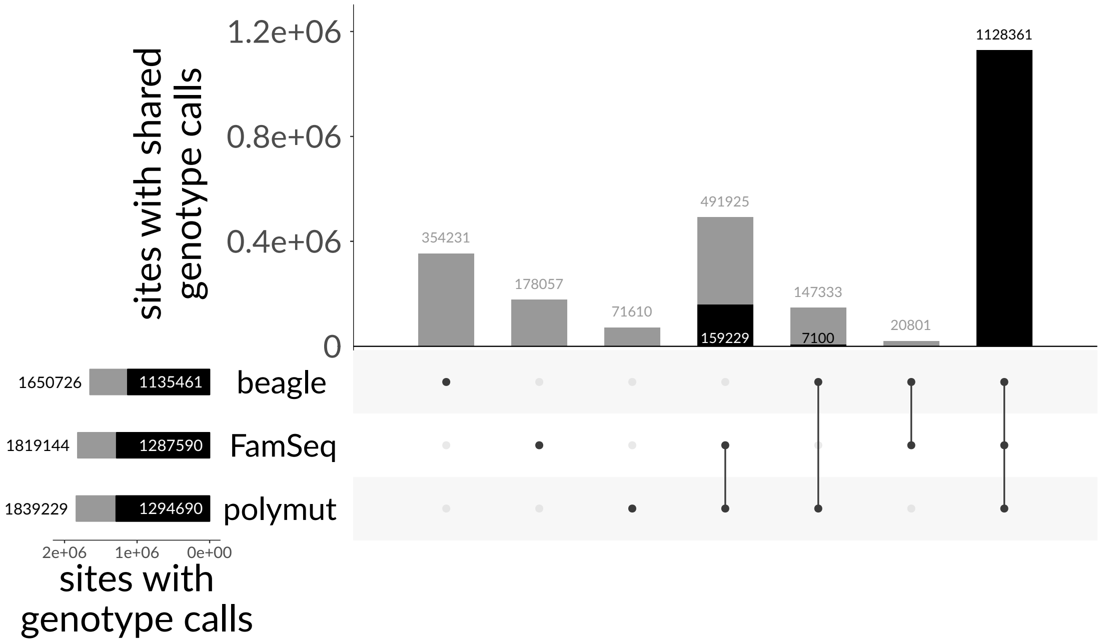

Figure S 4: Constructing the germline ground truth genotype for our granulocytes dataset from the consensus of three pedigree-aware callers. Variants were called on a previously generated whole exome bulk sequencing data set of the person's family (Figure 2), which is separate from the single cell and bulk whole exome sequencing data generated for this study. Variants were called using beagle, FamSeq and polymutt with total counts of individual tools (bottom left bar plot) and tool intersections of sites called (top bar plot) in grey. Black fractions of bars represent the sites selected for our consensus: sites where either all callers agree upon a genotype or at least two callers agree upon a particular genotype and the third tool has a missing genotype (i.e. doesn't contradict the genotype called by the other two).

PNG for our own dataset, see Figure 2). Candidate sites to call were generated using bcftools mpileup, taking all sites where at least one read covered a non-reference nucleotide in the single cell or its respective bulk. We ran two different modes of the software: Calling sites with a minimum read coverage of 1 in the single cell sample (default mode in Figure 3) or including zero-coverage sites, effectively imputing genotypes at these sites from the bulk sample (imputation). The different filter thresholds used represent the false discovery rate (`--fdr`) that we can control for (Section S 1.6).

For MonoVar (version `0.0.1`, which is commit `e0cf2db` cleaned up for the bioconda version we created), parameters were set as recommended in the software repository README [Zafar]. We ran two different modes of the software, with consensus filtering between cells of a population (`-c 1`) and without the filtering (`-c 0`). The different filter thresholds used represent the parameter `-t` being varied. However, turning on the consensus filtering did not cause any substantial changes, possibly due to the low number of cells sequenced (Figure S 5). Varying the parameter `-t` did not cause any substantial changes in MonoVar results, either.

SCAN-SNV [Luquette et al., 2019] was installed via the install instruction in the repository on 2019-10-30[11] and patched locally to make it run. The pipeline was then run with parameters set as recommended in the software repository `README.md` and the command line help, with the only changes made for a better comparability with the other tools. Namely, we tried to call as sensitively as possible by setting: `--min-sc-alt 1 --min-sc-dp 1 --min-bulk-dp 1`. However, the default pipeline only provides calls for somatic variants (variants only found in single cells, not in the bulk, as opposed to

[11] This are the install instructions at the repository state at `https://github.com/parklab/scan-snv/blob/62b7dcbc21e6cac64dc384abdfa708120e33819c/README.md`. This includes the following packages and versions from the custom conda channel `https://anaconda.org/jluquette`: `r-scansnv 0.1`, `scansnv 0.9`, `r-fastghquad 1.0` and `natefoo-slurm-drmaa 1.2.0`. Further, it requires manual registration of the java executable for `gatk 3.8`.

all the present alternative alleles), heavily filtering candidate sites before calling. We thus attempted a fairer alternative allele calling comparison by modifying the pipeline to remove the filter that eliminates alternative alleles also seen in the bulk sample and the filter that eliminates known dbsnp variation[12]. The different filter thresholds used represent the `--fdr` parameter, which—according to the authors—does not formally control the false discovery rate [Luquette et al., 2019].

For SCcaller (version `1.2`, as available on bioconda), default parameters were set as recommended in the software repository README [Dong, 2018]. We ran four different modes of the software, varying two things: 1. For the heterozygous candidate sites that SCcaller requires as input, we either used dbSNP [Sherry et al., 2001, NCBI, 2016] entries or generated them from the bulk background sample that was also used for calling in ProSolo, running GATK HaplotypeCaller. As the dbSNP-derived heterozygous candidate sites consistently outperformed those generate from the bulk background sample, only results using dbSNP are reported in the figures, to provide a cleaner comparison. 2. In addition to the recommended settings that require a minimum coverage of 10 (default, `--min 10 --minvar 4 --RD 20`), we also ran SCcaller with reduced coverage requirements to increase sensitivity for a fairer comparison with ProSolo (sensitive, `--min 4 --minvar 1 --RD 10`), thus including sites with a minimum read coverage of 4 and down to one read covering the alternative nucleotide. The three filtering thresholds used represent the filtering levels available for the `-a cutoff` command-line parameter. The filtering threshold gave certain leverage over the precision-recall trade-off and reducing the coverage requirements improved the sensitivity of results (Figure S 5).

For SCIPhI (version `0.1.4`, as available on bioconda), we ran two different modes: 1. The default parameters of the command line tool (`default`). 2. Fewer iterations

[12] I.e. we edited the snakemake `rule scansnv_somatic_sites` to exclude the following requirements: `tab[,bulk.alt] == 0 & tab[,bulk.idx] == '0/0' & tab$dbsnp == '.'`.

(`-l 400000`, because of the excessive run time) and more `sensitive` by turning off all heuristic filters (`--cwm 1 --mnp 1 --ms 1 --bns 0 --bnc 0 --ncf 0 --mnc 1`). On the whole exome dataset with five cells, SCIPhI ran for three weeks for each of these modes. On the whole genome dataset with two cells, an initial run of SCIPhI was killed due to a server problem after running for more than eight weeks. An attempt to restart from intermediate data using the `--il` command line parameter had the model estimations escalating towards unrealistic values, and the crash was thus left unrecoverable. A rerun on a server with more performant CPUs led to the sensitive mode finishing after 5 weeks and the default mode after 7.5 weeks. As no option for filtering or thresholding based on sensitivity vs. specificity is available, only two unfiltered data points are given in the respective plots (Figure S 5).

##### S 2.3 Alternative Allele Calling

The most precise single cell variant callers to date, SCcaller and SCIPhI, can only call the presence vs. the absence of an alternative allele (i.e. the heterozygous and the homozygous alternative genotypes called jointly). We have thus focused on this for the main part of our benchmarking.

For ProSolo, the posterior probability of the presence of an alternative allele is calculated with Equation 10, which represent the blue Event areas in Figures 1D and S 2.

Calculations of $p$recision and $r$ecall were based on true positives (TP, alternative allele present in the ground truth and called by the respective software), false positives (FP, the respective software calls the presence of an alternative allele that does not exist in the ground truth at that site), and false negatives (FN, the respective software makes no call or calls a homozygous reference genotype at a ground truth site with an alternative allele), using the formulas:

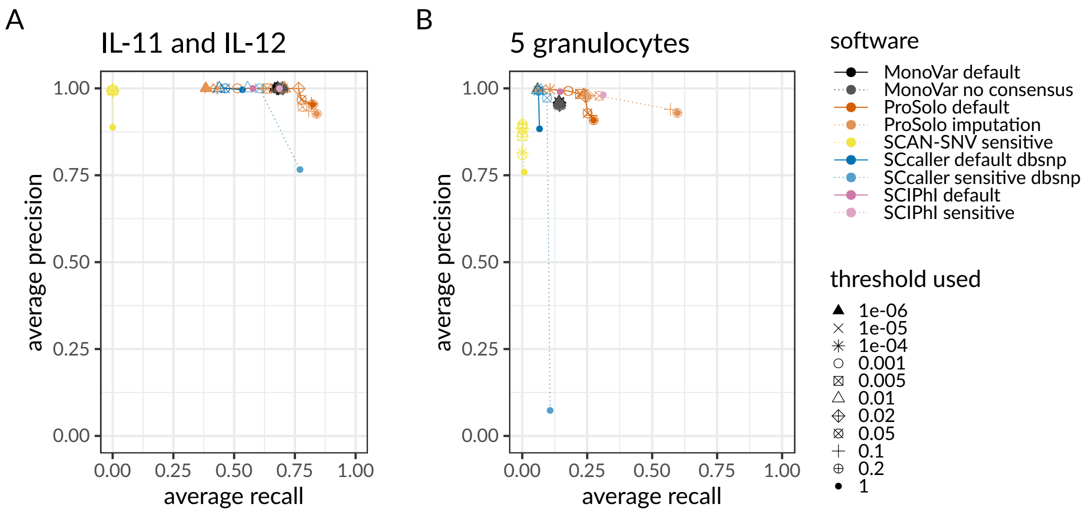

Figure S 5: Global view of the precision-recall plots for alternative allele calling across different software modes and filtering thresholds (for zoomed in view on the most interesting areas of the plot, see main manuscript Figure 3). **A** Precision and recall average of two whole genome sequenced single cells IL-11 and IL-12 against their kindred clone IL-1c as ground truth genotypes. **B** Precision and recall average of five whole exome sequenced single granulocytes against their pedigree-based germline genotype ground truth.
Thresholds used were applied via the following command line options: MonoVar `--t`; ProSolo `--fdr`; SCAN-SNV `--fdr`; SCcaller `-a cutoff`; SCIPhI `none available`. Software modes: MonoVar with consensus filtering (*default*) or without (*no consensus*); ProSolo with minimum coverage 1 in single cell (*default*), or imputing zero coverage sites based on bulk sample (*imputation*); SCcaller with recommended settings (*default*) or with a more *sensitive* calling; SCIPhI with default parameters (*default*) or all heuristics off (*sensitive*).

$$
\begin{aligned}
p &= \frac{TP}{TP + FP} \\
r &= \frac{TP}{TP + FN}
\end{aligned}
\tag{S 32}
$$

The whole genome cell line dataset (Figure 2A) seems much less challenging than the other dataset: all methods achieve high precision in alternative allele calling (Figures S 5A and 3A), at recall rates of 45% and higher. In comparison with all other tools, ProSolo achieves the most striking increases in recall of nearly 10%. For example, for a precision above 99%, its maximum recall is 76.6% compared to 70.5% for MonoVar,

68.7% for SCIPhI and 61.0% for SCcaller. SCAN-SNV achieves up to 99.2% precision, but only 0.01% recall. This can be explained by it aiming at somatic mutations, while the vast majority of SNVs in a genome will be germline variants.

Although a relative increase in recall of about 10% at utmost precision is certainly remarkable, ProSolo demonstrates its power on the second (whole exome) dataset. This dataset is considerably more challenging than the first one. See Figures 3B and S 5B for respective results. For this dataset, only SCcaller, SCIPhI and ProSolo achieved a precision above 99%, with ProSolo reaching a 20% increase of recall to 17.6%, compared to SCIPhI's 14.6%, and SCcaller with 7.2% (Figures 3B and S 5B). In comparison, MonoVar achieved a maximum precision of only 96.29%. However, this was at a much higher recall (14.13%) than for example SCcaller (9.55% at a precision of 97.23%). SCcaller's decreased recall on this dataset might be due to its estimation of local allelic bias by also taking biases at neighboring sites into account—in whole exome data the number of neighboring sites available for this estimation will be limited and might lead to less reliable estimates. On this dataset, SCAN-SNV's recall increased to 0.16% at a decreased maximum precision of 89.7%. Most likely, this decreased precision is an artefact of using the germline genotype as ground truth. At the sites with somatic mutations in single cells, which SCAN-SNV focuses on, this ground truth will instead contain the germline genotype and will incorrectly classify alternative alleles as false positives. Due to this effect, we also expect the calculated precision of all the other tools to be an underestimate, but against the backdrop of large numbers of sites where the single cells harbour the germline genotype, the effect will be much smaller. The same ground truth caveat also applies—inversely—for the recall of SCIPhI's sensitive mode and ProSolo's imputation mode. Whenever coverage of a site is missing in a single cell, SCIPhI may impute the genotype with the last common ancestor genotype of the most closely related cells, while ProSolo will impute to the majority genotype in the bulk sample. While both strategies provide a biologically meaningful imputation that will be more useful than post-hoc modes of imputation, we expect that a number of imputed sites were incorrectly classified as true positives with this germline ground truth. As this will lead to an overestimation of the recall, we have excluded both SCIPhI's sensitive mode and ProSolo's imputation mode from the discussion of this dataset—however, their results are nevertheless displayed in all respective figures.

ProSolo's improvements in accuracy can be further formalised using the $F_1$ score, the harmonic mean of precision $p$ and recall $r$, with $\beta = 1$ for the general $F$ score defined as [van Rijsbergen, 1979, Chinchor, 1992]:

$$F_\beta = (\beta^2 + 1) \times \frac{pr}{\beta^2 p + r} \tag{S 33}$$

The per-cell $F_1$ score clearly demonstrates ProSolo's performance advantage (Figure S 6), even when restricting our comparison to ProSolo's (and SCIPhI's) default mode for a fairer comparison on the whole exome dataset with a germline ground truth (see caveat discussed above).

##### S 2.4 False Discovery Rate Control of Alternative Allele Calling

Finally, a feature where ProSolo clearly stands out, is the control over the false discovery rate. As can be seen in Figure 3 (and Figure S 5), ProSolo provides flexible control over precision vs. recall via specifying a false discovery rate of interest. While no other tool provides a formal control over the easily interpretable false discovery rate, several of the tools provide other types of thresholds which we varied in attempts to achieve higher precision or recall. However, none of them provide control over similar ranges of precision and recall. Thus, ProSolo is the only tool that provides the user with the choice of either aiming for more discoveries at the cost of a higher rate of false discoveries, or at aiming for a more limited number of discoveries with higher confidence in each of them.

Further, Figure S 7A demonstrates, that ProSolo provides a rigorous control over the

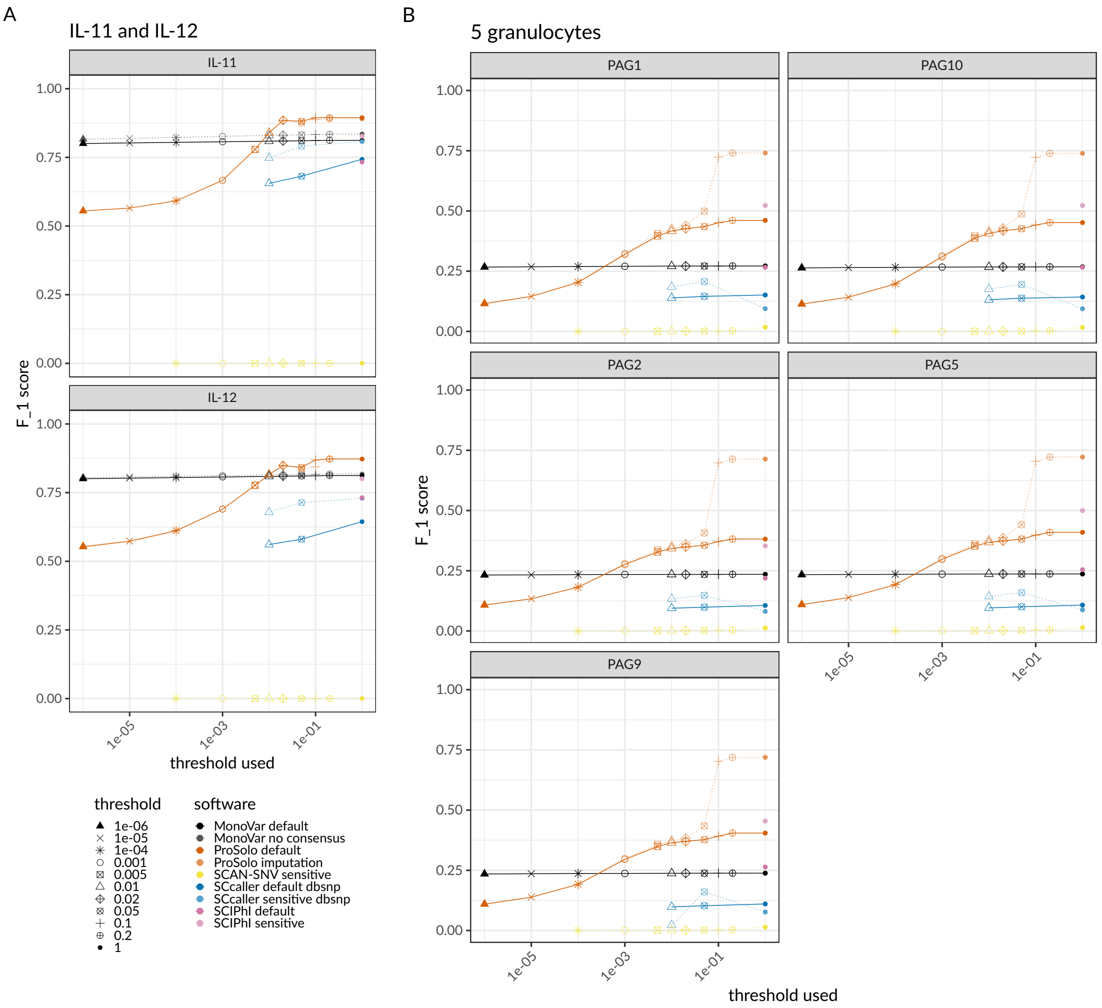

Figure S 6: Cell-specific $F_1$ scores for different filtering threshold levels for **A** the whole genome dataset and **B** the whole exome dataset.

Thresholds used were applied via the following command line options: MonoVar `--t`; ProSolo `--fdr`; SCAN-SNV `--fdr`; SCcaller `-a cutoff`; SCIPhI `none available`. Software modes: MonoVar with consensus filtering (*default*) or without (*no consensus*); ProSolo with minimum coverage 1 in single cell (*default*), or imputing zero coverage sites based on bulk sample (*imputation*); SCcaller with recommended settings (*default*) or with a more *sensitive* calling; SCIPhI with default parameters (*default*) or all heuristics off (*sensitive*).

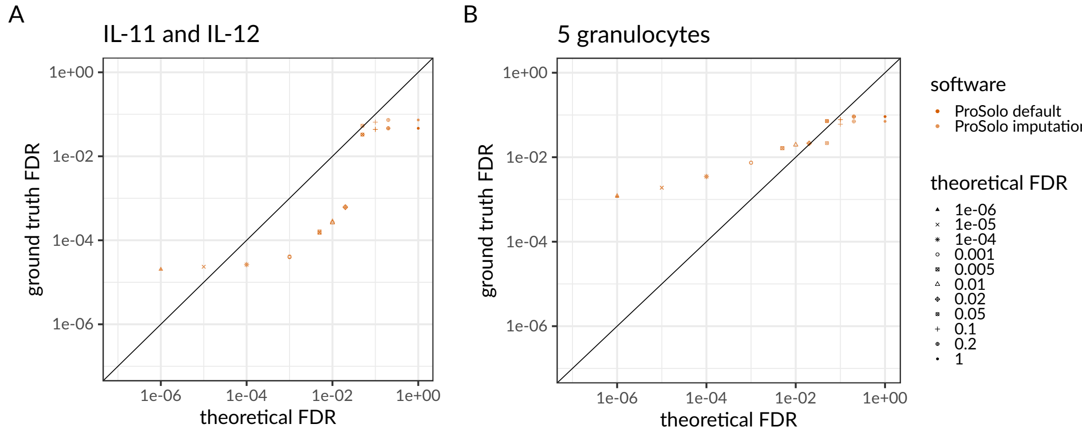

Figure S 7: False discovery rate (FDR, $1 - precision$) as calculated based on the ground truth, compared to the theoretical FDR we controlled for using ProSolo, for (A) the whole genome data set and (B) the whole exome data set. The diagonal indicates identity of the ground truth FDR with the FDR that was controlled for. Any points underneath it indicate overly conservative FDR control, any points above it indicate an FDR control that is too lenient.
Thresholds used were applied via the following command line options: MonoVar `--t`; ProSolo `--fdr`; SCAN-SNV `--fdr`; SCcaller `-a cutoff`; SCIPhI `none available`. Software modes: MonoVar with consensus filtering (*default*) or without (*no consensus*); ProSolo with minimum coverage 1 in single cell (*default*), or imputing zero coverage sites based on bulk sample (*imputation*); SCcaller with recommended settings (*default*) or with a more *sensitive* calling; SCIPhI with default parameters (*default*) or all heuristics off (*sensitive*).

false discovery rate on the whole genome dataset. The only exception—in both default and imputation mode—is when attempting to control for a false discovery rate below 0.0001. This indicates, that ProSolo's model still needs refinement in its resolution for sites with a very high likelihood of an alternative allele present. A similar trend can be seen on the whole exome dataset (Figure S 7B), but for any attempt to control for a false discovery rate below 0.02. This is within the common range of values often chosen as a false discovery rate to control for in practice and not being able to actually achieve the false discovery rate control at such values is usually unacceptable. However, as also discussed above, we generally expect the precision (of all tools) to be underestimated on this dataset: using the germline genotypes as ground truth results in misclassifying actual somatic variants in single cells as false positives. This underestimation of precision on this dataset in turn means an overestimation of the false discovery rate ($1-p$recision), and is thus to be expected. This artefact of using a germline ground truth also shows in the discrepancy in the ground-truth based false discovery rate between ProSolo's default and imputation mode when controlling for higher false discovery rates. As expected, imputation turned out less precise in the whole genome dataset (Figure S 7A). In contrast, imputation seems to have improved precision for the whole exome dataset (Figure S 7B). This would suggest that the larger call set from ProSolo's imputation mode (with a higher recall) is also more precise than the smaller call set from the default mode. However, this increase in precision is most likely explained by an imputation towards the majority genotype in the bulk sample that will usually favor the germline genotype at a site, no matter what the actual single cell genotype would be. Taken together, this indicates that the false discovery rate that we can calculate with the given germline ground truth for the whole exome dataset is higher than the actual value would be with a more accurate (somatic) ground truth available.

##### S 2.5 Allele dropout rate calculations

Leveraging our ground truths and using three different ways to calculate the allele dropout rate, we can confirm the general validity of the respective event definitions (Figure 1D) and evaluate limitations. For this, we will focus on the set of sites where the respective ground truth call is heterozygous, as these are the sites where the dropout of one of the alleles can be identified clearly.

The first way of calculating the allele dropout rate leverages the posterior probabilities generated by the ProSolo model (Equation S 31). At each ground truth heterozygous site, we sum up the probabilities of the two allele dropout events defined in ProSolo ("ADO to alt" and "ADO to ref" in Figure 1D) to obtain a total allele dropout probability (see Equation S 31). Obtaining the expected value of the allele dropout rate then constitutes adding those totals across all ground truth heterozygous sites with coverage and dividing by their number. This gives us an expected allele dropout rate based on the data and the ProSolo model ("expected value" in Figure 4):

$$\frac{\sum\limits_{\substack{\text{covered ground truth}\\ \text{heterozygous sites}}} P(\text{allele dropout} \mid \boldsymbol{Z}^s, \boldsymbol{Z}^b)}{\#\text{ covered ground truth heterozygous sites}} \tag{S 34}$$

The two ways to validate this expected allele dropout rate both depend on a comparison against the respective ground truth genotypes. One important difference in the ground truth used here, compared to the ground truth used for alternative allele calling, is that we further filtered the ground truth for the Dong et al. [2017] dataset. For this dataset, the basic ground truth was generated using GATK HaplotypeCaller and variant quality score recalibration. However, this recalibration optimises on the presence (heterozgyous or homozygous alternative) vs. the absence (homozygous reference) of an alternative allele and filtered genotypes will not reliably distinguish between heterozygous genotypes and homozygous alternative genotypes. We thus excluded all sites where the genotype quality (`GQ` FORMAT field) is below a PHRED score of 30. This corresponds to a 0.1% probability of having erroneously called the current genotype over the second most likely genotype, and thus also filters for a clear distinguishability between heterozygous and homozygous alternative genotypes. This is important both for identifying allele dropout and for identifying exact genotypes as in Section S 2.6.

The first validation of these rates from within ProSolo is to genotype all the ground truth heterozygous sites with ProSolo. We take the most likely genotype (i.e. the genotype whose coloured event area in Figure 1D has the highest joint probability). Then, every ground truth heterozygous site that ProSolo calls as homozygous is counted as a dropout site and the number of dropout sites is divided by the total number of ground truth heterozygous sites where the respective single cell had coverage. This gives an allele dropout rate based on a comparison against the ground truth ("hom at ground truth het" in Figure 4):

$$\frac{\#\text{ ground truth heterozygous sites called homozygous}}{\#\text{ covered ground truth heterozygous sites}} \tag{S 35}$$

A second validation of these rates is another comparison against the ground truth, but without using ProSolo in any way. For each single cell, using samtools mpileup, we identify all heterozygous ground truth sites with a coverage of at least 7. Without amplification bias we could be reasonably sure to sample both alleles of a heterozygous site at coverage 7 (the probability to sample only one allele with a coverage of 7 would be: $2 \cdot 0.5^7 = 0.015625$). We then count a site as an allele dropout, if there is one allele (reference or alternative) with no read coverage at all. The allele dropout rate is the fraction of allele dropout sites within the respective cell's heterozygous ground truth sites ("no alt/ref read at ground truth het" in Figure 4):

$$\frac{\text{\# ground truth heterozygous site with only one allele covered}}{\text{\# covered ground truth heterozygous sites}} \tag{S 36}$$

The expected allele dropout rates based on the ProSolo probabilities for allele dropout fall into the range of previously published allele dropout rates [Wang et al., 2014, Hou et al., 2012, Xu et al., 2012, Ling et al., 2009, Spits et al., 2006, Renwick et al., 2006, Lodato et al., 2015] ("published" in Figure 4). This analysis also shows that the ProSolo expected allele dropout rates, based on the model's probabilities, correspond with those based on the comparison of ProSolo genotypes with the ground truth (Figure 4), demonstrating that the explicit modelling of allele dropout events is useful for genotyping. However, in that comparison, the expected allele dropout rate is consistently underestimated on our own whole exome data ("granulocytes" in Figure 4), while it is slightly overestimated for the data from Dong et al. [2017] ("Dong 2017" in Figure 4). These deviations in the estimation in different directions suggest that the empirical distributions we currently use do not fit all datasets. Instead, they most likely depend on the respective whole genome amplification and library preparation procedures, as well as other possible batch effects. This conclusion is further supported by the much lower naively calculated allele dropout rate in the high coverage cells of the two datasets (IL-11 in Dong et al. [2017], PAG1 and PAG10 in our granulocytes, compare S 3). Here, the empirical distributions used seem to overestimate allele dropout and the performance of ProSolo could be further improved with distributions tailored to the specific dropout profile of the experiment on that individual cell. However, this overestimation of the allele dropout probability does not seem to impact the genotyping resolution (see below, Figures S 8 and S 9).

##### S 2.6 Genotype calling

As a further performance comparison, we want to include a direct comparison of the actual called genotypes against the ground truth genotypes of the respective data set. As only ProSolo, MonoVar and SCAN-SNV provide actual genotype calls—a further refinement over the alternative allele calling described above—SCcaller and SCIPhI cannot be included in this comparison. Further, the small numbers of calls made by SCAN-SNV would not be interpretable in the figures provided, and we have thus also excluded SCAN-SNV from this comparison.

Comparisons provided here are based on the software modes that achieved the highest $F_1$ score across most single cells for alternative allele calling (Figure 3C). For ProSolo, this amounts to the default mode without imputation (which also ensures a fair comparison against the germline-based ground truth) and controlling for a false discovery rate of 0.2. For MonoVar, this means no consensus filtering and no filtering via the `--t` thresholding parameter.

Plotting the overall precision and recall of ProSolo and MonoVar genotyping (Figure S 8), demonstrates that the higher recall of ProSolo is limited to the calling of heterozygous genotypes for the whole genome datasets (IL-11, IL-12 in Figure S 8A). For the whole exome dataset, ProSolo achieves a higher recall of both heterozygous and homozygous alternative ground truth genotypes (PAG cells in Figure S 8A). However, the recall remains much lower in this dataset, possibly due to the more uneven coverage of the respective cells and the lower coverage of the background bulk sample (PNG in Figure S 3).

Wherever ProSolo generates more genotype calls—be they heterozygous or homozygous alternative calls—it also makes more errors (Figure S 8B). In the only case where ProSolo does not generate more calls than MonoVar—homozygous alternative calls in the whole genome datasets—it also does not show a noticeable increase in false genotype

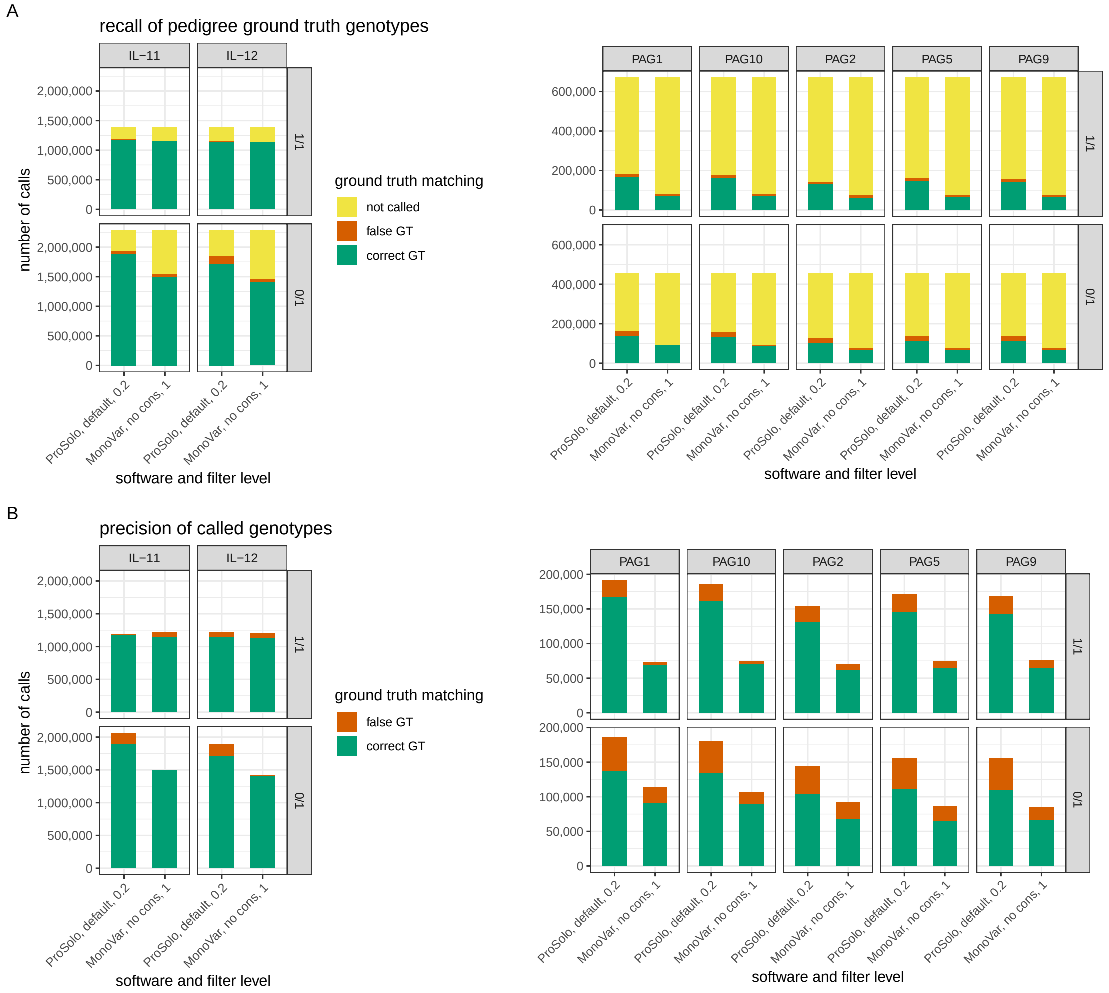

Figure S 8: Genotype calling precision and recall in comparison against the respective ground truth genotypes (from IL-1c in the left plots; from PNG in the plots on the right). MonoVar and ProSolo were run with the parameters that maximised the $F_1$ score alternative allele calling (Figure 3C). Facet labels at the top of plots indicate the single cell. Facet labels on the right side of plots indicate the genotype: "1/1" for homozgyous alternative, "0/1" for heterozygous. **A** Recall of ground truth genotypes. Here, facet labels on the right side of plots indicate the ground truth genotype compared against. ProSolo provides a genotype call at more of the ground truth sites, with a slight but noticeable increase of false genotypes provided at ground truth heterozygous sites. **B** Precision of called genotypes. Here, facet labels on the right side of plots indicate the genotype of the caller. The higher amount of calls by ProSolo (most noticeable for heterozygous calls for the whole genome datasets, and homozygous alternative calls for the whole exome datasets) trades off with a loss in precision.

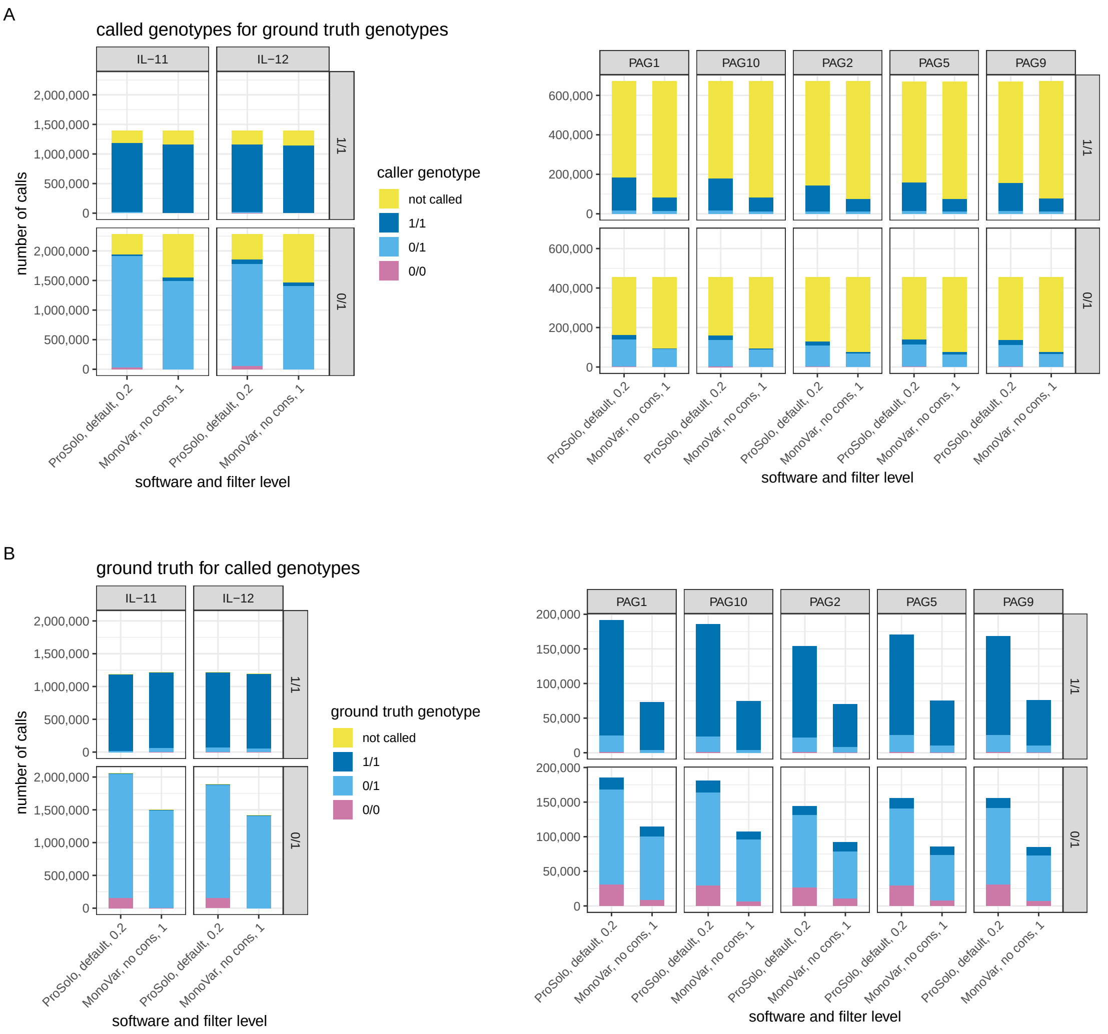

Figure S 9: Genotype calling comparison against the respective ground truth genotypes (from IL-1c in the left plots; from PNG in the plots on the right). MonoVar and ProSolo were run with the parameters that maximised the $F_1$ score alternative allele calling (Figure 3C). Panels A and B correspond to the respective panels in Figure S 8, but with a finer resolution of the false genotypes. Facets at the top of plots indicate the single cell. **A** Genotypes called by the respective tool are indicated by the colouring, for the respective ground truth genotypes as indicated by the facets on the right side of the plots. **B** Ground truth genotypes are indicated by the colouring, for the genotypes called by the respective tool as indicated by the facets on the right side of the plots. Both tools show visible amounts of misclassification in most of the possible categories.

calls.

When looking at the exact genotype miscalls (Figure S 9), it becomes apparent that the higher amount of miscalls at higher recall in ProSolo is biased for heterozygous calls made by ProSolo (Figure S 9B): Heterozygous calls made by ProSolo are more often at ground truth homozygous reference sites, whereas heterozygous calls at ground truth homozygous alternative sites do not occur more often than for MonoVar. Thus, while MonoVar heterozygous calls are more likely to be miscalled homozygous alternative sites than miscalled homozygous reference sites, the inverse holds for ProSolo. This indicates, that both models are not yet well-balanced. For ProSolo, a further improvement could be to adapt the amplification bias distributions currently used (see Section S 1.2.2 for some suggestions). Also, learning distributions from the data at hand could further improve precision and recall wherever data is more uniformly distributed than in the dataset the current distributions are derived from [Lodato et al., 2015].

Altogether, the genotype-level analysis also confirms that a higher and more uniform coverage of the single cell data leads to higher precision and recall in the variant calling, no matter which caller is used. E.g. for the—most evenly covered—whole genome sequencing data (Figure S 3A), both callers achieve a very high recall of ground truth homozygous alternative sites with a very high precision (left panels in Figures S 8 and S 9). Also, for the whole exome data (right panels in Figures S 8 and S 9), recall is visibly higher for all genotypes across both tools in the samples with a higher and more uniform coverage (PAG1 and PAG10, see Figure S 3), and precision is noticeably higher in those samples for MonoVar.

##### S 2.7 Further Software and Resources used

All analysis code for both the main processing pipeline and the manuscript figures is available at: `https://github.com/prosolo/benchmarking_prosolo` (or as preserved by Zenodo at: `https://doi.org/10.5281/zenodo.3769116`). An overview and cita-

Table S 3: Software and related resources used in this manuscript. All software versions as installed with conda.

| software / resource | version | purpose | citation |
|---|---|---|---|
| cutadapt | 1.14 | adapter trimming | [Martin, 2011] |
| trimmomatic | 0.36 | quality trimming | [Bolger et al., 2014] |
| bwa mem | 0.7.15 | read alignment / mapping | [Li, 2013] |
| samtools | 1.4.1 | handle SAM / BAM files | [Li et al., 2009] |
| picardtools | 2.9.2 | mark duplicates | |
| bcftools | 1.6 | handle VCF / BCF files | [Danecek et al., 2011] |
| mosdepth | 0.2.5 | coverage calculations | [Pedersen and Quinlan, 2018] |
| Okabe Ito palette | | colorblind-safe | [Okabe and Ito, 2008] |
| ggthemes | 0.2.5 | Okabe Ito palette | [Arnold et al., 2019] |
| Brewer palettes | | colorblind-friendly | [Brewer, 2006] |
| RColorBrewer | 1.1_2 | Brewer palettes | [Neuwirth, 2014] |
| tidyverse | 1.1.1<br>1.3.0 | ground truth comparisons<br>results data handling | [Wickham et al., 2019] |
| ggplot2 | 3.2.1 | result plots | [Wickham, 2016] |
| Inkscape | 0.92.4 | info-graphics, improved labels | |
| VGAM | 1.1_1 | distributions in R | [Yee, 2008, 2015] |
| UpsetR | | set intersection plot S 4 | [Conway et al., 2017] |

tions of software and resources beyond those mentioned in the methods section are provided in Table S 3.

### References

J. B. Arnold, G. Daroczi, B. Werth, B. Weitzner, J. Kunst, B. Auguie, B. Rudis, H. W. C. f. t. g. package.), J. T. C. f. t. l. package), and J. London. ggthemes: Extra

Themes, Scales and Geoms for 'ggplot2', May 2019. URL https://CRAN.R-project.org/package=ggthemes.

Y. Benjamini and Y. Hochberg. Controlling the False Discovery Rate: A Practical and Powerful Approach to Multiple Testing. *Journal of the Royal Statistical Society. Series B (Methodological)*, 57(1):289–300, 1995. URL http://www.jstor.org/stable/2346101.

A. M. Bolger, M. Lohse, and B. Usadel. Trimmomatic: a flexible trimmer for Illumina sequence data. *Bioinformatics*, 30(15):2114–2120, Aug. 2014. ISSN 1367-4803. doi: 10.1093/bioinformatics/btu170. URL https://academic.oup.com/bioinformatics/article/30/15/2114/2390096. Publisher: Oxford Academic.

C. Brewer. ColorBrewer: Color Advice for Maps, 2006. URL http://colorbrewer2.org/#.

B. L. Browning and S. R. Browning. Improving the Accuracy and Efficiency of Identity-by-Descent Detection in Population Data. *Genetics*, 194(2):459–471, June 2013. ISSN 0016-6731, 1943-2631. doi: 10.1534/genetics.113.150029. URL http://www.genetics.org/content/194/2/459.

N. Chinchor. MUC-4 Evaluation Metrics. In *Proceedings of the 4th Conference on Message Understanding*, MUC4 '92, pages 22–29, Stroudsburg, PA, USA, 1992. Association for Computational Linguistics. ISBN 978-1-55860-273-1. doi: 10.3115/1072064.1072067. URL https://doi.org/10.3115/1072064.1072067. event-place: McLean, Virginia.

J. R. Conway, A. Lex, and N. Gehlenborg. UpSetR: an R package for the visualization of intersecting sets and their properties. *Bioinformatics*, 33(18):2938–2940, Sept. 2017. ISSN 1367-4803. doi: 10.1093/bioinformatics/btx364. URL https://academic.oup.com/bioinformatics/article/33/18/2938/3884387.

P. Danecek, A. Auton, G. Abecasis, C. A. Albers, E. Banks, M. A. DePristo, R. E. Handsaker, G. Lunter, G. T. Marth, S. T. Sherry, G. McVean, and R. Durbin. The variant call format and VCFtools. *Bioinformatics*, 27(15):2156–2158, Aug. 2011. ISSN 1367-4803. doi: 10.1093/bioinformatics/btr330. URL `https://academic.oup.com/bioinformatics/article/27/15/2156/402296`. Publisher: Oxford Academic.

X. Dong. SCcaller README (https://github.com/biosinodx/SCcaller/tree/61629e8c39a5b9c0e0891bdff7a9 Aug. 2018. URL `https://github.com/biosinodx/SCcaller/tree/61629e8c39a5b9c0e0891bdff7a9785f5028beec`. original-date: 2016-06-09T15:43:47Z.

X. Dong, L. Zhang, B. Milholland, M. Lee, A. Y. Maslov, T. Wang, and J. Vijg. Accurate identification of single-nucleotide variants in whole-genome-amplified single cells. *Nature Methods*, 14(5):491–493, Mar. 2017. ISSN 1548-7091, 1548-7105. doi: 10.1038/nmeth.4227. URL `http://www.nature.com/doifinder/10.1038/nmeth.4227`.

F. Eggenberger and G. Pólya. Über die Statistik verketteter Vorgänge. *ZAMM - Journal of Applied Mathematics and Mechanics / Zeitschrift für Angewandte Mathematik und Mechanik*, 3(4):279–289, 1923. ISSN 1521-4001. doi: 10.1002/zamm.19230030407. URL `https://onlinelibrary.wiley.com/doi/abs/10.1002/zamm.19230030407`.

B. Grüning, R. Dale, A. Sjödin, B. A. Chapman, J. Rowe, C. H. Tomkins-Tinch, R. Valieris, and J. Köster. Bioconda: sustainable and comprehensive software distribution for the life sciences. *Nature Methods*, 15(7):475–476, July 2018. ISSN 1548-7105. doi: 10.1038/s41592-018-0046-7. URL `https://www.nature.com/articles/s41592-018-0046-7`.

J. Hoell, M. Gombert, S. Ginzel, S. Loth, P. Landgraf, V. Käfer, M. Streiter, A. Prokop, M. Weiss, R. Thiele, and A. Borkhardt. Constitutional Mismatch Repair-deficiency and Whole-exome Sequencing as the Means of the Rapid Detection of the Causative

MSH6 Defect. *Klinische Pädiatrie*, 226(06/07):357–361, Nov. 2014. ISSN 0300-8630, 1439-3824. doi: 10.1055/s-0034-1389905. URL http://www.thieme-connect.de/DOI/DOI?10.1055/s-0034-1389905.

Y. Hou, L. Song, P. Zhu, B. Zhang, Y. Tao, X. Xu, F. Li, K. Wu, J. Liang, D. Shao, and e. al. Single-Cell Exome Sequencing and Monoclonal Evolution of a JAK2\mbox-Negative Myeloproliferative Neoplasm. *Cell*, 148(5):873–885, Mar. 2012. doi: 10.1016/j.cell.2012.02.028. URL http://dx.doi.org/10.1016/j.cell.2012.02.028. bibtex: Hou2012.

htslib. C library for high-throughput sequencing data formats. URL https://github.com/samtools/htslib.

J. O. Irwin. A Distribution Arising in the Study of Infectious Diseases. *Biometrika*, 41(1/2):266–268, 1954. ISSN 0006-3444. doi: 10.2307/2333023. URL https://www.jstor.org/stable/2333023.

N. L. Johnson, A. W. Kemp, and S. Kotz. *Univariate discrete distributions.* Wiley, Hoboken, N.J, 3rd ed edition, 2005. ISBN 978-0-471-27246-5.

J. Köster. Rust-Bio: a fast and safe bioinformatics library. *Bioinformatics*, 32(3):444–446, Feb. 2016. ISSN 1367-4803, 1460-2059. doi: 10.1093/bioinformatics/btv573. URL http://bioinformatics.oxfordjournals.org/content/32/3/444.

J. Köster, L. Dijkstra, T. Marschall, and A. Schönhuth. Enhancing sensitivity and controlling false discovery rate in somatic indel discovery. *bioRxiv*, page 741256, Aug. 2019. doi: 10.1101/741256. URL https://www.biorxiv.org/content/10.1101/741256v1.

B. Li, W. Chen, X. Zhan, F. Busonero, S. Sanna, C. Sidore, F. Cucca, H. M. Kang, and G. R. Abecasis. A Likelihood-Based Framework for Variant Calling and De Novo

Mutation Detection in Families. *PLoS Genet*, 8(10):e1002944, Oct. 2012. doi: 10.1371/journal.pgen.1002944. URL http://dx.doi.org/10.1371/journal.pgen.1002944.

H. Li. Aligning sequence reads, clone sequences and assembly contigs with BWA-MEM. *arXiv:1303.3997 [q-bio]*, Mar. 2013. URL http://arxiv.org/abs/1303.3997. arXiv: 1303.3997.

H. Li, B. Handsaker, A. Wysoker, T. Fennell, J. Ruan, N. Homer, G. Marth, G. Abecasis, and R. Durbin. The Sequence Alignment/Map format and SAMtools. *Bioinformatics*, 25(16):2078–2079, Aug. 2009. ISSN 1367-4803. doi: 10.1093/bioinformatics/btp352. URL https://academic.oup.com/bioinformatics/article/25/16/2078/204688. Publisher: Oxford Academic.

J. Ling, G. Zhuang, B. Tazon-Vega, C. Zhang, B. Cao, Z. Rosenwaks, and K. Xu. Evaluation of genome coverage and fidelity of multiple displacement amplification from single cells by SNP array. *MHR: Basic science of reproductive medicine*, 15(11): 739–747, Nov. 2009. ISSN 1360-9947. doi: 10.1093/molehr/gap066. URL https://academic.oup.com/molehr/article/15/11/739/1000481.

M. A. Lodato, M. B. Woodworth, S. Lee, G. D. Evrony, B. K. Mehta, A. Karger, S. Lee, T. W. Chittenden, A. M. D'Gama, X. Cai, L. J. Luquette, E. Lee, P. J. Park, and C. A. Walsh. Somatic mutation in single human neurons tracks developmental and transcriptional history. *Science*, 350(6256):94–98, Oct. 2015. ISSN 0036-8075, 1095-9203. doi: 10.1126/science.aab1785. URL http://science.sciencemag.org/content/350/6256/94.

L. J. Luquette, C. L. Bohrson, M. A. Sherman, and P. J. Park. Identification of somatic mutations in single cell DNA-seq using a spatial model of allelic imbalance. *Nature Communications*, 10(1):1–14, Aug. 2019. ISSN 2041-

1723. doi: 10.1038/s41467-019-11857-8. URL https://www.nature.com/articles/s41467-019-11857-8.

M. Martin. Cutadapt removes adapter sequences from high-throughput sequencing reads. *EMBnet.journal*, 17(1):10–12, May 2011. ISSN 2226-6089. doi: 10.14806/ej.17.1.200. URL https://journal.embnet.org/index.php/embnetjournal/article/view/200. Number: 1.

P. Müller, G. Parmigiani, C. Robert, and J. Rousseau. Optimal Sample Size for Multiple Testing: The Case of Gene Expression Microarrays. *Journal of the American Statistical Association*, 99(468):990–1001, Dec. 2004. ISSN 0162-1459, 1537-274X. doi: 10.1198/016214504000001646. URL http://www.tandfonline.com/doi/abs/10.1198/016214504000001646.

P. Müller, G. Parmigiani, and K. Rice. FDR and Bayesian multiple comparisons rules. *Proceedings of the ISBA 8th World Meeting on Bayesian Statistics (Valencia)*, June 2006. URL http://www.bepress.com/jhubiostat/paper115.

R. C. NCBI. Database resources of the National Center for Biotechnology Information. *Nucleic Acids Research*, 44(D1):D7–D19, Jan. 2016. ISSN 0305-1048. doi: 10.1093/nar/gkv1290. URL https://academic.oup.com/nar/article/44/D1/D7/2503096.

E. Neuwirth. RColorBrewer: ColorBrewer Palettes, Dec. 2014. URL https://cran.r-project.org/web/packages/RColorBrewer/index.html.

M. Okabe and K. Ito. Color Universal Design (CUD) / Colorblind Barrier Free, Sept. 2008. URL https://jfly.uni-koeln.de/color/.

B. S. Pedersen and A. R. Quinlan. Mosdepth: quick coverage calculation for genomes and exomes. *Bioinformatics*, 34(5):867–868, Mar. 2018. ISSN 1367-4803. doi: 10.1093/bioinformatics/btx699. URL https://academic.oup.com/bioinformatics/article/34/5/867/4583630. Publisher: Oxford Academic.

G. Peng, Y. Fan, T. B. Palculict, P. Shen, E. C. Ruteshouser, A.-K. Chi, R. W. Davis, V. Huff, C. Scharfe, and W. Wang. Rare variant detection using family-based sequencing analysis. *Proceedings of the National Academy of Sciences*, 110(10):3985–3990, Mar. 2013. ISSN 0027-8424, 1091-6490. doi: 10.1073/pnas.1222158110. URL `http://www.pnas.org/content/110/10/3985`.

G. Peng, Y. Fan, and W. Wang. FamSeq: A Variant Calling Program for Family-Based Sequencing Data Using Graphics Processing Units. *PLoS Comput Biol*, 10(10): e1003880, Oct. 2014. doi: 10.1371/journal.pcbi.1003880. URL `http://dx.doi.org/10.1371/journal.pcbi.1003880`.

Á. J. Picher, B. Budeus, O. Wafzig, C. Krüger, S. García-Gómez, M. I. Martínez-Jiménez, A. Díaz-Talavera, D. Weber, L. Blanco, and A. Schneider. TruePrime is a novel method for whole-genome amplification from single cells based on *Tth*PrimPol. *Nature Communications*, 7:13296, Nov. 2016. ISSN 2041-1723. doi: 10.1038/ncomms13296. URL `https://www.nature.com/articles/ncomms13296`.

P. J. Renwick, J. Trussler, E. Ostad-Saffari, H. Fassihi, C. Black, P. Braude, C. M. Ogilvie, and S. Abbs. Proof of principle and first cases using preimplantation genetic haplotyping – a paradigm shift for embryo diagnosis. *Reproductive BioMedicine Online*, 13(1):110–119, Jan. 2006. ISSN 1472-6483. doi: 10.1016/S1472-6483(10)62024-X. URL `http://www.sciencedirect.com/science/article/pii/S147264831062024X`.

rust htslib. HTSlib bindings and a high level Rust API for reading and writing BAM files. URL `https://github.com/rust-bio/rust-htslib`.

S. T. Sherry, M.-H. Ward, M. Kholodov, J. Baker, L. Phan, E. M. Smigielski, and K. Sirotkin. dbSNP: the NCBI database of genetic variation. *Nucleic Acids Research*,

29(1):308–311, Jan. 2001. ISSN 0305-1048. doi: 10.1093/nar/29.1.308. URL https://academic.oup.com/nar/article/29/1/308/1116004.

J. Singer, J. Kuipers, K. Jahn, and N. Beerenwinkel. Single-cell mutation identification via phylogenetic inference. *Nature communications*, 9(1):5144–5144, 2018. ISSN 2041-1723. doi: 10.1038/s41467-018-07627-7. URL https://europepmc.org/articles/PMC6279798/.

C. Spits, C. Le Caignec, M. De Rycke, L. Van Haute, A. Van Steirteghem, I. Liebaers, and K. Sermon. Whole-genome multiple displacement amplification from single cells. *Nature Protocols*, 1(4):1965–1970, Nov. 2006. ISSN 1750-2799. doi: 10.1038/nprot.2006.326. URL https://www.nature.com/articles/nprot.2006.326.

M. Taschuk and G. Wilson. Ten simple rules for making research software more robust. *PLOS Computational Biology*, 13(4):e1005412, Apr. 2017. ISSN 1553-7358. doi: 10.1371/journal.pcbi.1005412. URL http://journals.plos.org/ploscompbiol/article?id=10.1371/journal.pcbi.1005412.

C. van Rijsbergen. Evaluation. In *Information Retrieval*, pages 112–140. Butterworths, London, 2 edition, 1979. URL http://www.dcs.gla.ac.uk/Keith/Preface.html.

Y. Wang, J. Waters, M. L. Leung, A. Unruh, W. Roh, X. Shi, K. Chen, P. Scheet, S. Vattathil, H. Liang, A. Multani, H. Zhang, R. Zhao, F. Michor, F. Meric-Bernstam, and N. E. Navin. Clonal evolution in breast cancer revealed by single nucleus genome sequencing. *Nature*, 512(7513):155–160, Aug. 2014. ISSN 0028-0836. doi: 10.1038/nature13600. URL http://www.nature.com/nature/journal/v512/n7513/full/nature13600.html.

H. Wickham. *ggplot2: Elegant Graphics for Data Analysis*. Springer-Verlag New York, 2016. ISBN 978-3-319-24277-4. URL https://ggplot2.tidyverse.org.

H. Wickham, M. Averick, J. Bryan, W. Chang, L. D. McGowan, R. François, G. Grolemund, A. Hayes, L. Henry, J. Hester, M. Kuhn, T. L. Pedersen, E. Miller, S. M. Bache, K. Müller, J. Ooms, D. Robinson, D. P. Seidel, V. Spinu, K. Takahashi, D. Vaughan, C. Wilke, K. Woo, and H. Yutani. Welcome to the tidyverse. *Journal of Open Source Software*, 4(43):1686, 2019. doi: 10.21105/joss.01686.

X. Xu, Y. Hou, X. Yin, L. Bao, A. Tang, L. Song, F. Li, S. Tsang, K. Wu, H. Wu, and e. al. Single-Cell Exome Sequencing Reveals Single-Nucleotide Mutation Characteristics of a Kidney Tumor. *Cell*, 148(5):886–895, Mar. 2012. doi: 10.1016/j.cell.2012.02.025. URL `http://dx.doi.org/10.1016/j.cell.2012.02.025`. bibtex: Xu2012.

T. W. Yee. The VGAM Package. *R News*, 8(2):28–39, Oct. 2008. URL `http://CRAN.R-project.org/doc/Rnews/`.

T. W. Yee. *Vector Generalized Linear and Additive Models: With an Implementation in R*. Springer, New York, NY, USA, 2015.

H. Zafar. MonoVar README (https://bitbucket.org/hamimzafar/monovar/src/7b47571d85a377c425feea86[illegible] URL `https://bitbucket.org/hamimzafar/monovar/src/7b47571d85a377c425feea86f6e591ee832fe342`.

H. Zafar, Y. Wang, L. Nakhleh, N. Navin, and K. Chen. Monovar: single-nucleotide variant detection in single cells. *Nature Methods*, 13(6):505–507, June 2016. ISSN 1548-7105. doi: 10.1038/nmeth.3835. URL `https://www.nature.com/articles/nmeth.3835`.
